# Identification of septoria nodorum blotch susceptibility genes in hard winter wheat

**DOI:** 10.64898/2026.05.13.724689

**Authors:** Anju Maan Ara, Danielle J. Holmes, Timothy L. Friesen, Brett F. Carver, Guihua Bai, Paul St. Amand, Amy Bernado, Rajat Sharma, Meriem Aoun

## Abstract

*Key message* Characterized and unknown septoria nodorum blotch susceptibility/resistance genes were identified in contemporary U.S. hard winter wheat.

The necrotrophic fungus *Parastagonospora nodorum* is the causal agent of septoria nodorum blotch (SNB) of wheat. To determine the prevalence of SNB sensitivity genes in a contemporary U.S. hard winter wheat (HWW), we evaluated a panel of 619 breeding lines and cultivars against five *P. nodorum* isolates and five necrotrophic effectors (NEs), SnToxA, SnTox1, SnTox3, SnTox267 and SnTox5, and genotyped the panel using genotyping-by-sequencing (GBS) markers and diagnostic Kompetetive-allele specific PCR (KASP) markers for the sensitivity genes *Tsn1-B1*, *Snn1-B1*, and *Snn3-B1/B2*. GBS analysis identified 34,357 GBS-single nucleotide polymorphism (SNP) markers. Evaluations against *P. nodorum* isolates showed that 40-67% of the genotypes were susceptible in the panel. Toxin infiltration assays showed that 54%, 2%, 37%, 13%, and 15% of the genotypes were sensitive to SnToxA, SnTox1, SnTox3, SnTox267, and SnTox5, respectively. Diagnostic KASP markers for *Tsn1-B1*, *Snn1-B1*, and *Snn3-B1/B2* showed prediction accuracies of 98%, 75%, and 92% for the corresponding effectors SnToxA, SnTox1, and SnTox3, respectively. Genome-wide association studies (GWAS) not only confirmed the presence of the previously characterized sensitivity genes *Tsn1-B1*, *Snn1-B1*, *Snn2*, *Snn3-B1/B2*, and *Snn5-B1*, but also identified new loci to be associated with responses to *P. nodorum* isolates and NEs. Of which, *Qsnb.osu-2AS* on chromosome 2AS was associated with responses to all five isolates. We developed KASP markers *KASP_S4B_643615365*, *KASP_ S2D_16184991*, and *KASP_S2A_9833162* linked to *Snn5-B1*, *Snn2*, and *Qsnb.osu-2AS*, respectively. These findings should guide breeding for SNB resistance in hard winter wheat.

## Introduction

Common wheat (*Triticum aestivum* L., 2n = 6x = 42, AABBDD) is cultivated on more than 215 million hectares globally (Singh et al. 2022) and is one of the most important food crops. Hard winter wheat (HWW) is the most produced market class in the United States and is mostly cultivated in the Great Plains. However, wheat production is reduced due to multiple abiotic and biotic stresses. Septoria nodorum blotch (SNB), caused by the necrotrophic fungus, *Parastagonospora nodorum* (Berk.) (syn. *Septoria nodorum*, *Stagonospora nodorum*; teleomorph *Phaeosphaeria*), has become an economically important disease over the past several decades in many wheat-growing areas worldwide, including the eastern U.S., parts of northern Europe, Australia, and parts of North Asia (Bhathal et al. 2003; Murray and Brennan 2009; Crook et al. 2012; Figueroa et al. 2017; Cowger et al. 2020). *Parastagonospora nodorum* infects both leaves, causing leaf blotch, and spikes, causing glume blotch. This disease can reduce yield by up to 50% on susceptible cultivars (Liu et al. 2004b). In the Great Plains, SNB has become more prevalent in recent years (Aoun and Carver 2024). Farming practices such as minimum or no tillage that leave more crop residue on the soil surface, could have contributed to the increased prevalence of SNB. Furthermore, a study of *P. nodorum* isolates collected from multiple wheat growing regions in the U.S. showed that isolates collected from HWW in Oklahoma had higher genetic and virulence diversity (Richards et al. 2019). This diversity may have contributed to adaptation to HWW cultivars grown in Oklahoma.

Genetic dissection of wheat−*P. nodorum* pathosystem has revealed that the recognition of *P. nodorum* necrotrophic effectors (NEs) by host sensitivity genes in an inverse gene-for-gene manner leads to programed cell death in the host and consequently wheat susceptibility (Friesen and Faris 2021; Peters Haugrud et al. 2022; Kariyawasam et al. 2023). To date, seven wheat sensitivity genes/alleles (*Tsn1-B1*, *Snn1-B1*, *Snn2*, *Snn3-B1*, *Snn3-B2*, *Snn3-D1*, and *Snn5-B1*) (Running 2022; Seneviratne et al. 2024b, a; Zhang et al. 2025; Running et al. 2025) and five *P. nodorum* NE genes (*SnToxA*, *SnTox1*, *SnTox3*, *SnTox5*, and *SnTox267*) have been cloned (Ciuffetti et al. 1997; Liu et al. 2004a, 2012; Friesen et al. 2007, 2008, 2012; Gao et al. 2015; Shi et al. 2015a). Thirteen sensitivity wheat gene/allele-NE interactions have been identified, including *Tsn1-B1*–*SnToxA* (Friesen et al. 2006; Zhang et al. 2009; Friesen et al. 2009; Faris and Friesen 2009; Faris et al. 2010, 2011; Running et al. 2025), *Tsn1-B2*–*SnToxA* (Running et al. 2025), *Snn1-B1*–*SnTox1* (Liu et al. 2004a, b; Reddy et al. 2008; Liu et al. 2012; Shi et al. 2016a; Seneviratne et al. 2024a), *Snn1-B2*–*SnTox1* (Seneviratne et al. 2024a), *Snn2*–*SnTox267* (Friesen et al. 2007, 2009; Zhang et al. 2009; Richards et al. 2021), *Snn3-B1*–*SnTox3* (Friesen et al. 2008; Liu et al. 2009; Shi et al. 2016b; Zhang et al. 2025), *Snn3-B2*–*SnTox3* (Zhang et al. 2025), *Snn3-D1*–*SnTox3* (Friesen et al. 2008; Liu et al. 2009; Zhang et al. 2011), *Snn4*–*SnTox4* (Abeysekara et al. 2009, 2012), *Snn5*-*B1*–*SnTox5* (Friesen et al. 2012; Sharma et al. 2019; Running 2022; Kariyawasam et al. 2022), *Snn5*-*B2*–*SnTox5* (Running 2022), *Snn6*–*SnTox267* (Gao et al. 2015; Richards et al. 2022), and *Snn7*–*SnTox267* (Shi et al. 2015; Richards et al. 2022).

Although previous studies have identified SNB sensitivity genes and resistance loci and their prevalence in various wheat classes (Phan et al. 2018; Ruud et al. 2019; Cowger et al. 2020; Peters Haugrud et al. 2023; Szabo-Hever et al. 2025a), data specific to contemporary HWW grown in the U.S. Great Plains are limited. It has been reported that differences in *P*. *nodorum* sensitivity genes between wheat germplasm do exist (Crook et al. 2012; Richards et al. 2019). Given the rising incidence of SNB in the HWW production regions, elucidating the genetic basis of resistance and sensitivity in HWW is essential to guide effective selection within breeding programs. In this study, we investigated the genetics of susceptibility/resistance to SNB in a panel of 619 HWW genotypes, including elite breeding lines developed by the Oklahoma State University (OSU) wheat breeding program, along with cultivars from both the OSU and Kansas State University (KSU) breeding programs. We evaluated the panel at the seedling stage against *P. nodorum* isolates and NEs. Furthermore, we determined the prevalence of the sensitivity genes *Tsn1-B1*, *Snn1-B1*, *Snn3-B1*, and *Snn3-B2* in the panel using available diagnostic kompetitive allele specific PCR (KASP) markers (Seneviratne et al. 2024a; Zhang et al. 2025; Running et al. 2025). To identify genomic regions associated with SNB susceptibility/resistance in this panel, a genome-wide association study (GWAS) was conducted using the phenotypic data, single-nucleotide polymorphism (SNP) generated *via* genotyping-by-sequencing (GBS), and diagnostic KASP markers available for some characterized sensitivity genes. GWAS-significant SNPs associated with novel resistance/susceptibility loci, as well as characterized sensitivity genes that currently lack available diagnostic markers, were targeted for the development of KASP markers.

## Materials and Methods

### Plant materials and genotyping

A panel of 619 elite HWW breeding lines and cultivars was used in this study. This panel includes 532 doubled-haploid breeding lines selected from 14 bi-parental crosses and 38 OSU elite breeding lines from the OSU wheat breeding program, and 49 cultivars from the OSU and KSU wheat breeding programs (Supplementary Table S1). This panel was genotyped using GBS (Poland and Rife 2012) at the USDA-ARS Central Small Grain Genotyping Lab in Manhattan, KS. SNP genotype calling was performed using TASSEL software v.5 (Bradbury et al. 2007), and the physical positions of SNPs were assigned using the Chinese Spring RefSeqv2.1 developed by International Wheat Genome Consortium (IWGSC) (Zhu et al. 2021). SNP markers with missing data ≥ 60% were excluded, and missing data were imputed using Beagle 5 (Browning et al. 2018). Markers with minor allele frequency (MAF) ≤ 5%, and heterozygosity ≥ 15% were removed from further analysis, which left 34,357 SNP markers (Supplementary Table S2) for downstream analyses.

Additionally, the HWW panel was genotyped using diagnostic KASP markers available for three previously characterized sensitivity genes *Tsn1* (*Tsn1-B1_1Ka* and *Tsn1-B1_2Ka*), *Snn1* (*Snn1_null*), and *Snn3* (*Snn3-B1* and *Snn3-B2*), (Seneviratne et al. 2024; Zhang et al. 2025; and Running et al. 2025) (Supplementary Fig. S1). Genomic DNA was extracted from leaf tissue collected at the second leaf stage using a modified cetyl-trimethyl-ammonium-bromide (CTAB) method (Riede and Anderson 1996; Liu et al. 2006). DNA concentration was quantified using a Nanodrop spectrophotometer ND-8000 (ThermoFisher Scientific, Waltham, MA), and DNA stocks were diluted to a working concentration of 30 ng µL^−1^. KASP assay was performed following the modified LGC bioscience technologies thermal-cycling protocol (Makhoul et al. 2020). Each 100 µL primer mix contained 12 µL of each allele-specific primer (100 µM), 30 µL of the common primer (100 µM), and 46 µL of ddH_2_O. Each PCR reaction consisted of 5 µL of 2 × KASP master mix (KBS-1050-102; LGC Biosearch Technologies Hoddesdon, United Kingdom), 0.14 µL primer mix, 1.86 µL of ddH_2_O and 3 µL of DNA (30 ng/µL). The PCR was conducted on a Bio-Rad CFX-96 Opus real-time PCR system (Bio-Rad, Hercules, CA) with 94°C for 15 min, followed by 10 touchdown cycles of denaturation at 94°C for 20 s, annealing-extension starting at 65°C for 1 min with a decrement of 0.8°C at each cycle, followed by 30 cycles of denaturation at 94°C for 20 s, and annealing-extension at 57°C for 1 min. The fluorescent signal was measured after cooling down to 37°C. Endpoint fluorescence reading and genotype clustering were performed in Bio-Rad CFX Maestro Software 2.3. Up to three additional cycles of 94°C for 20 s and 57°C for 1 min were added to the previous protocol if the clusters were not well separated after initial PCR.

### Seedling evaluation using *P. nodorum* isolates

*Parastagonospora nodorum* isolates OKG16-Sn1, OKG16-Sn2, OKG16-Sn9, OKG16-Sn13, and OKG16-Sn16 used in this study were collected from the winter wheat cultivar ‘Gallagher’ in Canadian County, Oklahoma in 2016 (Richards et al. 2019). These isolates were separated based on their responses to the differential set of wheat lines (Supplementary Table S3) and in the presence/absence of effectors SnToxA, SnTox1, SnTox3, SnTox267, and SnTox5 (Table 1). The isolates were grown on V8-potato dextrose agar (PDA) (150 ml of V8 juice, 3 g of CaCO_3_, 10 g of Difco PDA, 10 g of agar in 1,000 ml of water) as described by Liu et al. (2004b). The cultures were grown for 12-14 days at 21°C with constant UVB light (Zilla Slimeline desert fixture 50UVB T8 Fluorescent bulb). After pycnidia sporulation, plates with fungal spores were washed with sterile distilled water and scraped with a sterile glass slide. The spore concentration was measured with a hemocytometer and adjusted to a final concentration of 1 × 10^6^ spores ml^−1^ using distilled water. Two drops of Tween 20 (polyoxyethylene sorbitan monolaurate) were added per 100 ml of inoculum to reduce the surface tension.

**Table 1.** Characterized necrotrophic effectors in five isolates of *Parastagonospora nodorum* and percentages of resistant and susceptible hard winter wheat genotypes to these isolates.

| Isolate ID | Present effectors | Absent effectors | Number of resistant genotypes | Number of intermediate genotypes | Number of susceptible genotypes |
| --- | --- | --- | --- | --- | --- |
| OKG16Sn-1 | SnToxA, SnTox1, SnTox267, and SnTox5 | SnTox3 | 126 (21%) | 195 (32%) | 286 (47%) |
| OKG16Sn-2 | SnToxA, SnTox1, SnTox267, SnTox3, and SnTox5 |  | 48 (8%) | 153 (25%) | 407 (67%) |
| OKG16Sn-9 | SnToxA, SnTox3, and SnTox5 | SnTox1 and SnTox267 | 55 (9%) | 308 (51%) | 242 (40%) |
| OKG16Sn-13 | SnToxA, SnTox1, SnTox267, and SnTox5 | SnTox3 | 53 (9%) | 298 (49%) | 255 (42%) |
| OKG16Sn-16 | SnToxA, SnTox1, SnTox267, SnTox3, and SnTox5 |  | 30 (5%) | 315 (52%) | 262 (43%) |

The 619 wheat genotypes were grown in the greenhouse at 20°C /18°C (day/night) with a 16-h photoperiod. The genotypes were planted in 72-cell trays filled with a commercial ‘Ready-Earth’ soil mix (Sun Gro). The susceptible check ‘Gallagher’ was planted as a border in each tray to reduce border effect. In each experiment, 4-5 seeds per genotype were planted in an augmented design and repeated 3-4 times per isolate (Supplementary Table S1). The differential lines TAM 110, Alsen, LP29, Jerry, Massey, ITMI38, F/G9519, Scoop1, Chinese Spring, BG220, and BG261 were included as checks alongside each experiment (Supplementary Table S3). Jack’s classic all-purpose (20-20-20) fertilizer was applied once a week to the seedlings according to the manufacturer’s instructions. At the second leaf stage, which is approximately 12-13 days post-planting, seedlings were uniformly inoculated using a hand-pressured sprayer until runoff, air-dried for 30 min, then placed in a mist chamber in the dark at 21°C with 100% relative humidity for 24 h. Thereafter, the plants were transferred to a growth chamber with a 12-h photoperiod, constant temperature of 21°C, and light intensity of 2,000 umol until disease evaluation. At seven days of post-inoculation, the plants were evaluated for their SNB reactions using a scale of 0-to-5 (Liu et al. 2004b), where 0 = absence of visible lesions (highly resistant); 1 = few penetration points, with lesions exhibiting flecking or small dark spots (resistant); 2 = lesions exhibiting dark spots with little surrounding necrosis or chlorosis (moderately resistant); 3 = dark lesions completely surrounded by necrosis or chlorosis and lesions of 2 to 3 mm (moderately susceptible); 4 = larger necrotic or chlorotic lesions of 4 mm or larger and with little coalescence (susceptible); and 5 = large coalescent lesions with very little remaining green leaf tissue (highly susceptible) (Liu et al. 2004b).

### Effector infiltration

We used five NEs, SnToxA, SnTox1, SnTox3, SnTox5, and SnTox267, produced by *P. nodorum.* SnToxA, SnTox1, and SnTox3 were expressed in *Pichia pastoris* as described in Friesen and Faris (2012). SnTox267 was expressed using a bacterial cell lysate method (Outram et al. 2021), whereas SnTox5 was isolated from a 3-week-old fungal culture filtrate from an avirulent *P. nodorum* strain transformed with *SnTox5* gene (Liu and Friesen 2012; Kariyawasam et al. 2024). The effector infiltration assays were performed at the USDA-ARS Cereal Crops Improvement Research Unit, Fargo, ND, USA. Infiltration was done following the protocol by Friesen and Faris (2012). Plants to be infiltrated were placed under fluorescent lights for 20 min to promote stomata opening. For each NE, two replicates per genotype were planted with two plants per replicate. If there was a discrepancy in the genotype reactions between the two replicates, a third replication was performed. The lines BG261, Chinese Spring, BG220, BG223, and LP29 were also planted alongside the toxin infiltration experiments as sensitive checks for SnToxA, SnTox1, SnTox3, SnTox267, and SnTox5, respectively (Supplementary Table S1). At the second leaf stage, approximately 20 μl of NE was infiltrated into the second leaf (one leaf per plant) using a needleless 1 ml syringe. The NE was infiltrated into the leaf by pressing the open end of the syringe into the leaf. The liquid was pushed intercellularly *via* the stomatal openings, leaving a water-soaked appearance on the leaves, marked with a permanent marker. After infiltration, the plants were placed in a growth chamber at 21°C and 12-h photoperiod. The reactions to NE effectors were scored at 3 days post-infiltration using a 0-to-3 scale, where 0 = no reaction (insensitive), 1 =slight but visible chlorosis, 2 = visible chlorosis/necrosis without tissue collapse, and 3 (highly sensitive) = necrosis with complete tissue collapse (Friesen and Faris 2012). For each NE assay, the susceptible check reaction was considered as a baseline to classify the genotypes into sensitive or insensitive.

Homogeneity of variances among experiments or replicates for each NE and *P. nodorum* isolate was assessed using Bartlett’s test in R (Aslam 2020). When Bartlett’s test indicated a violation of variance homogeneity, pairwise comparisons were performed using the Games-Howell test (Sauder and DeMars 2019). Only those experiments or replicates that exhibited homogeneous variances were retained for subsequent analyses.

### Population structure and linkage disequilibrium

Population structure and linkage disequilibrium (LD) in the wheat panel was performed as described by Lakkakula et al. (2025). Population structure was assessed using principal component analysis (PCA) and based on 34,357 GBS-SNP markers. PCA was performed with the ‘prcomp’ function in R and the population structure was visualized using the first two principal components (PCs). Linkage disequilibrium (LD) between SNP marker pairs was calculated in TASSEL version 5.2 (Bradbury et al. 2007) as the square of the correlation coefficient (*r^2^*). The function “geom_smooth” in the R package “ggplot2” was used to create a locally weighted estimated scatter plot smoother (LOESS) curve (Cleveland 1979) on the LD decay plot.

### Genome-wide association study

A genome wide association study (GWAS) was performed using phenotypic data, 34,357 filtered and imputed GBS-SNPs and diagnostic KASP markers for the sensitivity genes *Tsn1-B1*, *Snn1-B1*, *Snn3-B1*, and *Snn3-B2*. GWAS was conducted using three models: the Bayesian-information and linkage-disequilibrium iteratively nested keyway (BLINK) (Huang et al. 2019), the fixed and random model circulating probability unification (FarmCPU) (Liu et al. 2016), and the mixed linear model (MLM) (Yu et al. 2006) implemented in the Genomic Association and Prediction Integrated Tool v3 (GAPIT 3) (Wang and Zhang 2021) in the R software. The single-locus MLM is traditionally the most used model for GWAS. It controls spurious connections using kinship or family relatedness (K matrix) and population structure (Q matrix) (Zhang et al. 2005; VanRaden 2008). However, this model is likely to produce spurious associations because it was developed to evaluate one marker at a time (Wen et al. 2018). Multi-locus models such as FarmCPU and BLINK were later developed and are considered to be more accurate and reliable than single-locus models (Vikas et al. 2022; Sharma et al. 2025). The model FarmCPU works iteratively, using both fixed and random models, and includes significant markers as cofactors in each iteration to control for false associations without overfitting the model (Liu et al. 2016). The GWAS model BLINK is an upgraded version of FarmCPU that includes two key changes: 1) BLINK does not presume that the causal genes are evenly distributed throughout the genome, and 2) it excludes markers in LD with the most significant marker. To estimate maximum likelihood, BLINK uses the Bayesian information criterion (BIC) in a fixed-effect framework (Huang et al. 2019). In contrast to MLM, multi-locus models (BLINK and FarmCPU) identify distinct loci that are not in LD and tagged by their most significant markers. Previous studies reported that the BLINK model outperformed other GWAS models implemented in in GAPIT 3, including FarmCPU in terms of computing efficiency and statistical power, producing fewer false positives and detecting more real connections (Huang et al. 2019; Wang and Zhang 2021; Sharma et al. 2025).

The GWAS models used in this study incorporated the K matrix and the Q matrix based on PCA. The optimal number of principal components (PCs) included in the Q matrix was based on quantile–quantile (Q-Q) plots that illustrate the deviations between marker observed and expected -log10 (*P*) values (Aoun et al. 2021, 2022; Sharma et al. 2025). The number of PCs tested in the Q matrix was limited to the first four PCs. A threshold of false discovery rate (FDR) (Benjamini and Hochberg 1995) ≤ 0.05 was used to identify significant associations. The R package “CMplot” (https://github.com/YinLiLin/R-CMplot) and the “geom_point” function in the R package “ggplot2” (Wickham and Sievert 2009) were used to create the GWAS Manhattan plots.

### Development of KASP markers associated with response to septoria nodorum blotch

Major loci identified through GWAS that were linked to either unknown resistance/sensitivity genes or to sensitivity genes lacking reliable diagnostic markers in the literature were selected for KASP marker development, to facilitate their potential application in breeding programs. Twenty susceptible genotypes, carrying the susceptible allele of the marker, and 20 resistant genotypes, carrying the resistant allele of the marker, were selected from the GWAS panel to test the developed KASP markers in this study. The DNA of the samples was diluted to a concentration of 30 ng µL^−1^. In hexaploid wheat, homoeologous and paralogous regions can have substantial sequence similarity. Thus, locus-specific common primers were designed using the anchoring approach suggested by LGC Biosearch Technologies (https://biosearchassets.blob.core.windows.net/assetsv6/guide_kasp-assay-design-anchoring.pdf). A 200 bp sequence of the SNP of interest in the flanking region was obtained from the IWGSC RefSeq v2.1 (Zhu et al. 2021). This sequence was blasted to identify an anchor point (unique base) at the 3’ end to ensure locus-specific primers. HEX tail 5′-GAAGGTCGGAGTCAACGGATT-3′ and FAM tail 5′-GAAGGTGACCAAGTTCATGCT3′ were added to 5’ end of the SNP marker allele-specific primers. The designed primers were synthesized by Integrated DNA Technologies (Coralville, IA). The KASP reaction was performed using the modified LGC Biosearch Technologies protocol described by Makhoul et al. (2020). PCR reactions and conditions were described above in the method section ‘3.3.1 Plant materials and genotyping’.

## Results

### Wheat responses against *P. nodorum* isolates at the seedling stage

The reactions of 619 HWW genotypes to inoculation of the *P. nodorum* isolates ranged from 0 (highly resistant) to 5 (highly susceptible) (Fig. 1). The percentages of susceptible genotypes (having a disease reaction equal or higher than the susceptible checks) to isolates OKG16Sn-1, OKG16Sn-2, OKG16-Sn9, OKG16Sn-13, and OKG16Sn-16 were 47%, 67%, 40%, 42%, and 43%, respectively (Table 1). Genotypes with intermediate (moderate) reactions to the five isolates ranged from 25% (against isolate OKG16Sn-2) to 52% (against isolate OKG16Sn-16). There were low percentages of resistant genotypes to *P. nodorum* isolates, ranging from 5 –21% (Table 1). A total of 75 genotypes had resistant to intermediate reactions to all five isolates (Supplementary Table S4), including the OSU cultivars ‘Uncharted’, ‘Bentley’, ‘Big Country’, and ‘OK Corral’.

**Fig. 1.**
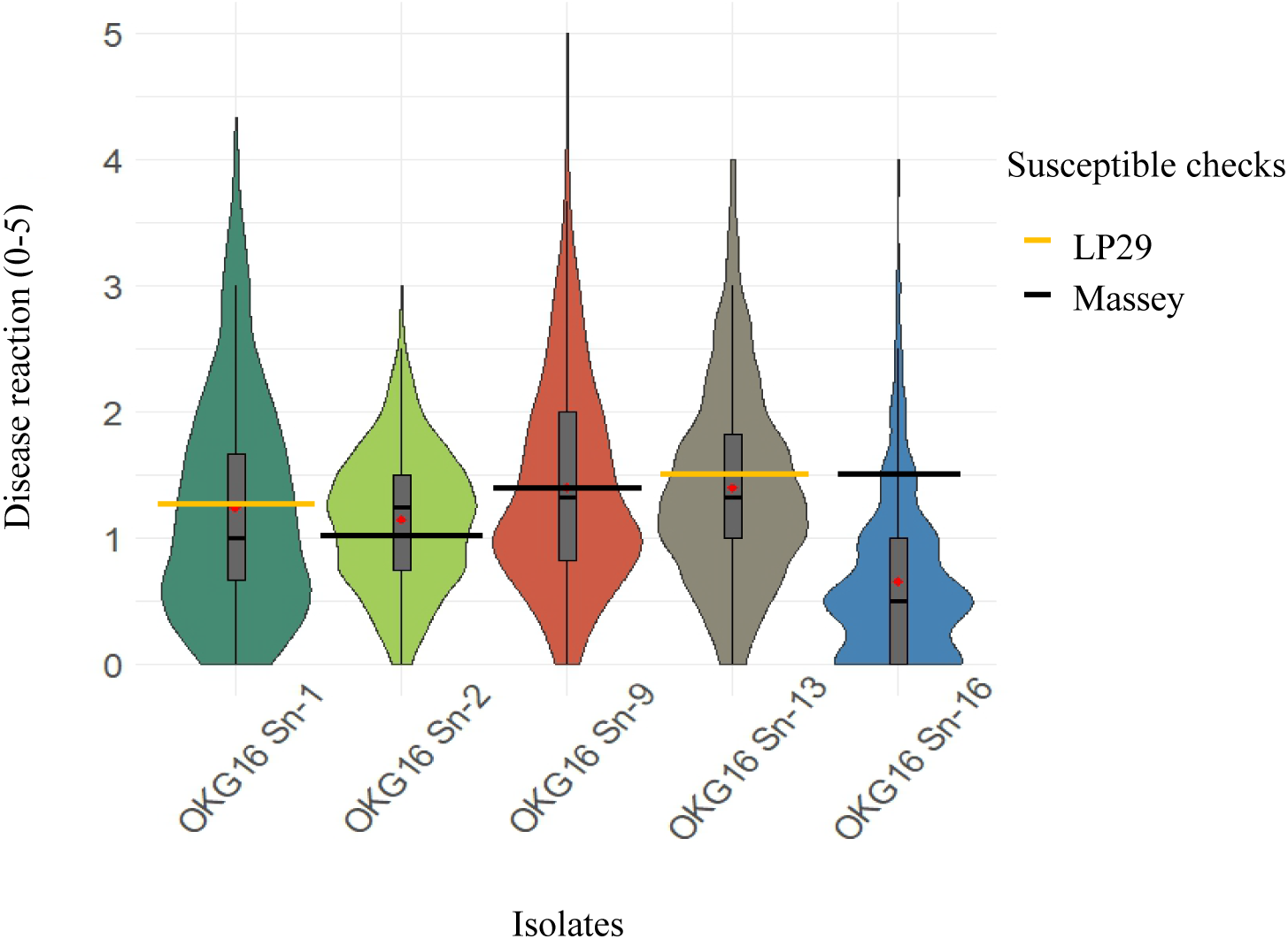
Distribution of disease reactions of 619 hard winter wheat genotypes against five *P. nodorum* isolates at the seedling stage. The red diamonds represent the mean disease reactions in this germplasm. The orange horizontal line indicates the reaction means of the susceptible check ‘LP29’ against isolates OKG16Sn-1 and OKG16Sn-13. The black horizontal line indicates the reaction means of the susceptible check ‘Massey’ against isolates OKG16Sn-2, OKG16Sn-9, and OKG16Sn-16.

### Reactions against *P. nodorum* effectors

Effector infiltration assays against five *P. nodorum* NEs showed that 54%, 2%, 37%, 13%, and 15% of the genotypes were sensitive to SnToxA, SnTox1, SnTox3, SnTox267, and SnTox5, respectively. Sensitive genotypes had reactions similar to those of the susceptible checks. The reaction means of the susceptible checks BG261 to SnToxA, Chinese Spring to SnTox1, BG220 to SnTox3, BG223 to SnTox267, and LP29 to SnTox5 were 3, 1.75, 2, 2, and 3, respectively (Fig. 2, Table 2). These results indicated that the sensitivity genes *Tsn1* and *Snn3* are the most frequent in this germplasm, whereas *Snn1* is the least frequent. In this HWW panel, we identified 153 insensitive genotypes to all five NEs (Supplementary Table S5), of which 37 were also resistant to the five tested *P. nodorum* isolates (Supplementary Table S6). This indicates the presence of additional effectors in *P. nodorum* isolates that are conditioning susceptibility in HWW. The OSU cultivars Big Country, Uncharted, and Bentley were found to be resistant against the five isolates and insensitive to all five tested NEs (Table 3, Supplementary Table S6). Although the cultivar ‘Green Hammer’ was insensitive to the five NEs, it was found to be susceptible to isolates OKG16Sn-1 and OKG16Sn-2 (Table 3), suggesting that these two isolates should carry other unknown effectors, conditioning susceptibility in Green Hammer.

**Fig. 2.**
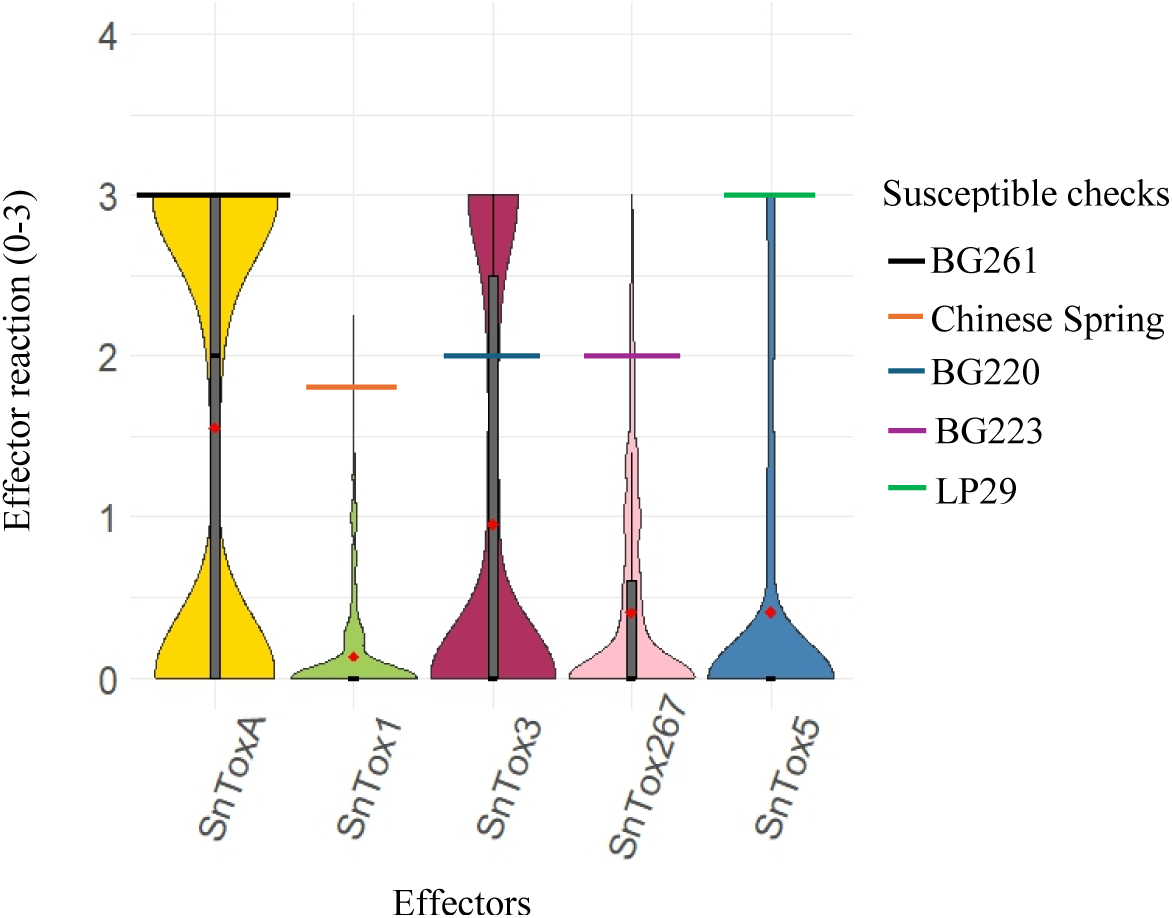
Distribution of reactions of 619 hard winter wheat to five *P. nodorum* effectors at the seedling stage. The red diamonds represent the mean reactions to effectors in this germplasm. The black, orange, blue, purple, and green horizontal lines indicate reaction means of the susceptible checks BG261, Chinese Spring, BG220, BG223, and LP29 to effectors SnToxA, SnTox1, SnTox3, SnTox267, and SnTox5, respectively.

**Table 2.**
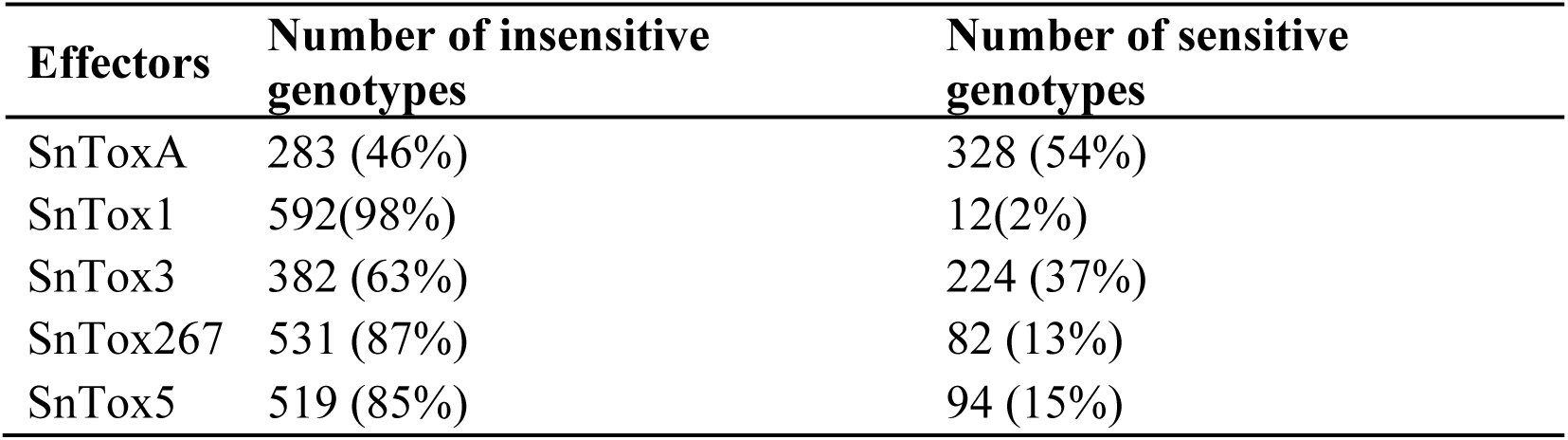
Percentages of insensitive and sensitive hard winter wheat genotypes to five *P. nodorum* effectors.

**Table 3.** Reactions of 32 hard winter wheat cultivars to *P. nodorum* isolates and effectors with associated sensitivity genes based on molecular markers.

| Cultivar Name | *P. nodourn* isolates <sup>a</sup> |  |  |  |  | *P. nodorum* effectors <sup>b</sup> |  |  |  |  | Sensitivity genes based on diagnostic markers <sup>c</sup> |  |  |  |  |
| --- | --- | --- | --- | --- | --- | --- | --- | --- | --- | --- | --- | --- | --- | --- | --- |
|  | OKG16Sn-1 | OKG16Sn-2 | OKG16Sn-9 | OKG16Sn-13 | OKG16Sn-16 | SnToxA | SnTox1 | SnTox3 | SnTox267 | SnTox5 | *Tsn1_1Ka* | *Tsn1_2Ka* | *Snn1_null* | *Snn3-B1* | *Snn3-B2* |
| Bentley | R | R | R | R | R | I | I | I | I | I | *Tsn1*- | *Tsn1*- | *Snn1*- | *Snn3-B1*- | *Snn3-B2*- |
| Big Country | R | R | R | M | M | I | I | I | I | I | *Tsn1*- | *Tsn1*- | *Snn1*- | *Snn3-B1*- | *Snn3-B2*- |
| Breakthrough | M | S | S | M | M | S | S | S | I | S | *Tsn1*+ | *Tsn1*+ | *Snn1*+ | *Snn3-B1*+ | *Snn3-B2*- |
| Butler's Gold | R | S | M | S | M | S | I | S | I | S | *Tsn1*+ | *Tsn1*+ | *Snn1*- | *Snn3-B1*+ | *Snn3-B2*- |
| Chisholm | S | S | M | S | S | S | I | S | S | I | *Tsn1*+ | *Tsn1*+ | *Snn1*- | *Snn3-B1*+ | *Snn3-B2*- |
| Double Stop CL+ | S | S | S | S | M | S | I | I | I | S | *Tsn1*+ | *Tsn1*+ | *Snn1*- | *Snn3-B1*- | *Snn3-B2*- |
| Duster | S | S | S | S | S | S | I | I | S | I | *Tsn1*+ | *Tsn1*+ | *Snn1*- | *Snn3-B1*- | *Snn3-B2*- |
| Firebox | M | S | M | M | M | S | I | I | I | I | *Tsn1*+ | *Tsn1*+ | *Snn1*- | *Snn3-B1*- | *Snn3-B2*- |
| Fuller | S | M | M | M | M | S | I | I | I | S | *Tsn1*+ | *Tsn1*+ | *Snn1*- | *Snn3-B1*- | *Snn3-B2*- |
| Gallagher | S | S | S | S | S | S | I | I | S | I | *Tsn1*+ | *Tsn1*+ | *Snn1*- | NA | *Snn3-B2*+ |
| Green Hammer | S | S | R | M | M | I | I | I | I | I | *Tsn1*- | *Tsn1*- | Het | *Snn3-B1*- | *Snn3-B2*- |
| High Cotton | S | S | M | S | M | S | I | I | I | I | *Tsn1*+ | *Tsn1*+ | *Snn1*+ | *Snn3-B1*- | *Snn3-B2*- |
| Iba | S | S | S | S | S | S | I | I | S | I | *Tsn1*+ | *Tsn1*+ | *Snn1*- | *Snn3-B1*+ | *Snn3-B2*- |
| Jagalene | M | S | M | M | M | S | I | S | I | I | *Tsn1*+ | *Tsn1*+ | *Snn1*- | *Snn3-B1*+ | *Snn3-B2*- |
| Jagger | S | S | S | S | M | S | I | I | I | S | *Tsn1*+ | *Tsn1*+ | Het | *Snn3-B1*- | *Snn3-B2*- |
| Karl | M | S | S | S | S | I | I | S | S | S | *Tsn1*- | *Tsn1*- | *Snn1*- | *Snn3-B1*+ | *Snn3-B2*- |
| Karl 92 | M | S | M | S | S | S | I | I | S | I | *Tsn1*- | *Tsn1*+ | *Snn1*- | *Snn3-B1*+ | *Snn3-B2*- |
| Kharkof | S | S | S | S | S | S | I | I | I | I | *Tsn1*+ | *Tsn1*+ | Het | *Snn3-B1*+ | *Snn3-B2*+ |
| Newton | S | M | S | S | M | S | I | I | I | S | *Tsn1*+ | *Tsn1*+ | *Snn1*- | *Snn3-B1*+ | *Snn3-B2*- |
| Nicoma | S | S | S | S | S | S | I | S | I | S | *Tsn1*+ | *Tsn1*+ | *Snn1*- | *Snn3-B1*+ | *Snn3-B2*+ |
| OK Bullet | S | S | M | S | M | S | I | S | I | S | *Tsn1*+ | *Tsn1*+ | Het | *Snn3-B1*+ | *Snn3-B2*- |
| OK Corral | R | M | M | R | M | I | I | S | I | I | *Tsn1*- | *Tsn1*- | *Snn1*- | *Snn3-B1*+ | *Snn3-B2*+ |
| Orange Blossom CL+ | R | S | M | S | M | I | I | S | I | I | *Tsn1*- | *Tsn1*- | *Snn1*- | *Snn3-B1*+ | *Snn3-B2*- |
| Overley | M | S | R | S | S | I | I | I | I | S | *Tsn1*- | *Tsn1*- | *Snn1*- | *Snn3-B1*- | *Snn3-B2*- |
| Paradox | S | S | S | S | S | S | S | I | S | I | *Tsn1*+ | *Tsn1*+ | *Snn1*+ | *Snn3-B1*+ | *Snn3-B2*+ |
| Ruby Lee | S | S | S | S | M | S | I | I | I | I | *Tsn1*+ | *Tsn1*+ | *Snn1*- | *Snn3-B1*+ | *Snn3-B2*+ |
| Showdown | M | R | M | M | S | S | I | I | I | I | *Tsn1*+ | *Tsn1*+ | *Snn1*- | *Snn3-B1*- | *Snn3-B2*- |
| Smith's Gold | M | R | R | S | S | S | I | S | I | I | *Tsn1*+ | *Tsn1*+ | Het | *Snn3-B1*+ | *Snn3-B2*- |
| Spirit Rider | M | S | M | M | S | I | I | I | S | I | *Tsn1*- | *Tsn1*- | Het | *Snn3-B1*- | *Snn3-B2*- |
| Triumph | M | S | S | S | S | S | I | I | S | I | *Tsn1*+ | *Tsn1*+ | *Snn1*- | *Snn3-B1*+ | *Snn3-B2*- |
| Turkey | M | S | S | S | M | S | I | I | S | I | *Tsn1*+ | *Tsn1*+ | Het | *Snn3-B1*+ | *Snn3-B2*+ |
| Uncharted | R | R | R | M | M | I | I | I | I | I | *Tsn1*- | *Tsn1*- | *Snn1*- | *Snn3-B1*- | *Snn3-B2*- |

<sup>a</sup> S= Susceptible, M=Moderate (or intermediate reactions), R=Resistant, NA= data not available due to germination issue <sup>b</sup> S=Sensitive, I=Insensitive <sup>c</sup> The sensitivity gene is present (+), the sensitivity gene is absent (-), NA: data not available, Het = Heterozygous call

### Frequencies of the sensitivity genes *Tsn1-B1*, *Snn1*, *Snn3-B1*, *Snn3-B2* based on KASP markers

Genotyping data for KASP markers (*Tsn1-B1_1Ka* and *Tsn1-B1_2Ka*) associated with *Tsn1-B1* showed that 328 genotypes (54%) carry the *Tsn1-B1* gene, and 280 genotypes (46%) lack this gene. Comparing *Tsn1-B1* marker results with SnToxA infiltration assay data, showed that several genotypes carrying *Tsn1-B1* based on markers (n= 5 genotypes for marker *Tsn1-B1_1Ka* and n=7 genotypes for marker *Tsn1_B1_2Ka*) but were insensitive to SnToxA, suggesting 2% and 3% of false positives for markers *Tsn1-B1_1Ka* and *Tsn1_B1_2Ka*, respectively. Markers revealed that 22% of the genotypes carry *Snn1*-B1, whereas 44% and 24% of the genotypes carry *Snn3-B1* and *Snn3-B2*, respectively. False positive rates for *Snn1-B1* and *Snn3-B1/Snn3-B2* markers were 25% (n=149 genotypes) and 12% (n=44 genotypes), respectively. Extremely low false negatives (genotypes carrying the sensitivity genes based on markers but insensitive to the corresponding NEs) were identified with these markers; 2% (n=5 genotypes) and 1% (n=4 genotypes) for *Tsn1* markers *Tsn1-B1_1Ka* and *Tsn1-B1_2Ka*, respectively, 0% for *Snn1-B1* marker, and 3% (n=6 genotypes) for *Snn3-B1/Snn3-B2* markers. The overall accuracy of markers *Tsn1-B1_1Ka*, *Tsn1-B1_2Ka*, *Snn1-B1*, and *Snn3-B1/Snn3-B2* were 98%, 98 %, 75%, and 92%, respectively.

### Linkage disequilibrium and PCA

Of the filtered 34,357 SNPs, 11,878 (34.6%) were mapped to the A genome, 16,150 (47%) to the B genome, 5,884 (17.1%) to the D genome, and 445 (1.3%) were unaligned (UN) to a chromosome. SNP numbers ranged from 322 on chromosome 4D to 3,685on chromosome 2D. The highest density of SNP markers in the 1 Mb window was observed on the A genome, whereas the lowest density of SNP markers was observed on the D genome (Supplementary Fig. S2). The genome-wise LD dropped to an *r*^2^ of 0.25 within 2.0 Mb on average (Supplementary Fig. S3). LD decayed to 0.25 at ∼1.2 Mb on average for genome A, at 2.5 Mb on average for genome B, and at 6.8 Mb on average for genome D (Supplementary Fig. S4). PCA using 34,357 SNPs showed low to moderate structure in the 619 HWW genotypes, where the first two PCs, PC1 and PC2, explained 10.7% and 6.0% of the genetic variation, respectively (Supplementary Fig. S5). The first 10 PCs accounted cumulatively for 35.5% of the variation.

### GWAS model selection

The most suitable GWAS model for each trait was selected based on the examination of the Q-Q plots. For all traits, the K matrix was included in the GWAS models, and the optimal number of PCs in the Q matrix varied by trait (Supplementary Fig. S6, Fig. S7). In the MLM model, the Q matrix was excluded (no PCs) for all traits except responses to isolates OKG16Sn-2 and OKG16Sn-9, where the first four PCs were included. For FarmCPU model, the first two PCs were included in the Q matrix for responses to all isolates except OKG16Sn-16, for which no PCs were included. Similarly, no PCs were included for responses to all NEs except SnTox267, for which the first two PCs were incorporated in the Q matrix of the model. In the BLINK model, no PCs were included for responses to isolates OKG16Sn-1, OKG16Sn-9, OKG16Sn-13, and OKG16Sn-16, as well as to the NEs SnToxA, SnTox1, and SnTox5, while the Q matrix included the first two PCs for responses to the NEs SnTox3 and SnTox267, and the first four PCs for response to isolate OKG16Sn-2.

### Markers associated with responses to *P. nodorum* isolates and necrotrophic effectors

For this study, we mainly reported results from the single-locus model MLM (Fig. 3 and 4, Supplementary Table S7) and the multi-locus model BLINK (Table 4, Supplementary Fig. S8 and S9). Based on the BLINK model, there were 23 significant loci associated with responses to the *P. nodorum* isolates, and 49 loci associated with responses to the NEs (Table 4). The highest number of significant SNPs associated with responses to *P. nodorum* isolates were positioned on chromosome arm 5BL, which corresponds to the position of *Tsn1-B1* gene (Table 4, Fig. 3, Supplementary Table S7, Supplementary Fig. S8). This indicates the importance of this gene in explaining susceptibility to these isolates, which all carry the NE SnToxA. Another locus on chromosome arm 2AS and designated here as *Qsnb.osu-2AS* was found to be important in explaining responses to all isolates. *Qsnb.osu-2AS* effects on responses to isolates OKG16Sn-2, OKG16Sn-13, and OKG16Sn-16 exceed that of the *Tsn1* locus (Table 4). This locus was not reported to be associated with any known SNB sensitivity/resistance genes. In addition to the two major loci on chromosome arms 5BL and 2AS, the BLINK model identified other unknown loci associated with responses to isolates OKG16Sn-2, OKG16Sn-9, OKG16Sn-13, and OKG16Sn-16. Unknown loci associated with SNB responses are on chromosome arms 1AS, 2BS, 2DS, 3BL, 5AS, 5AL, 6BS, 6BL, 6DS, 7BL (Table 4, Supplementary Fig. S8). GWAS identified the sensitivity genes *Tsn1-B1* (on chromosome 5BL), *Snn1-B1* (on chromosome 1BS), *Snn3-B1/Snn3-B2* (on chromosome 5BS), *Snn2* (on chromosome 2DS), *Snn5-B1* (on chromosome 4BL) to be associated with responses to the NE effectors SnToxA, SnTox1, SnTox3, SnTox267, and SnTox5, respectively (Table 4, Fig. 4). In addition to these genes, the BLINK model identified other unknown loci associated with responses to SnTox1, SnTox3, SnTox267, and SnTox5 were positioned on chromosome arms 1AS, 1AL, 1BL, 1DS, 2AL, 2BL, 3AS, 3AL, 3BS, 3BL, 4AL, 4BL, 5AS, 5AL, 6AL, 6BS, 6BL, 7BS, 7BL, and 7DL (Table 4, Supplementary Fig. S9).

**Fig. 3.**
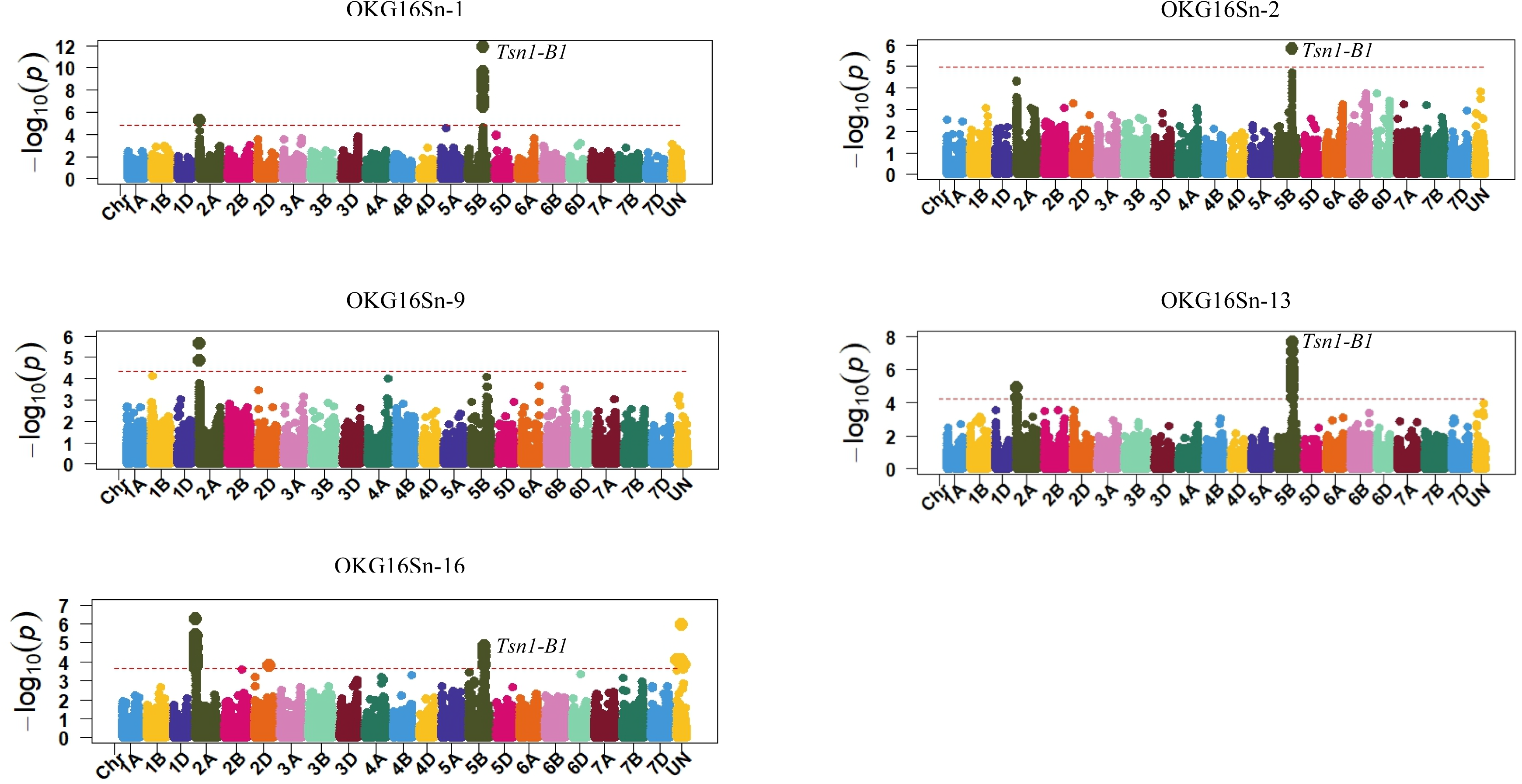
Manhattan plots showing significant markers associated with responses to five *P. nodorum* isolates using the MLM model. The horizontal red line indicates significance levels at a false discovery rate ≤ 0.05.

**Fig. 4.**
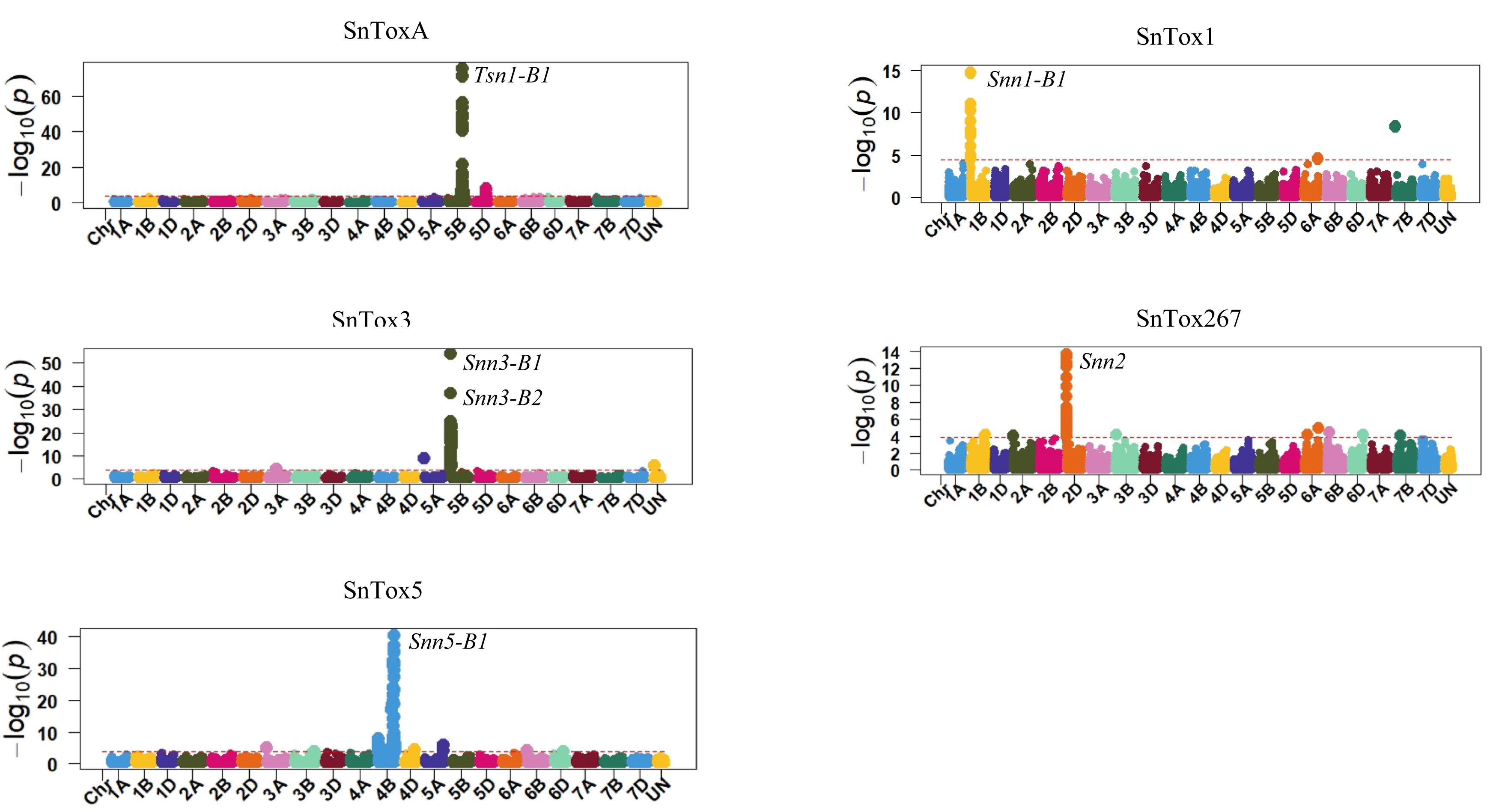
Manhattan plots showing significant markers associated with responses to five *P. nodorum* effectors using the MLM model. The horizontal red line indicates significance levels at a false discovery rate ≤ 0.05.

**Table 4.**
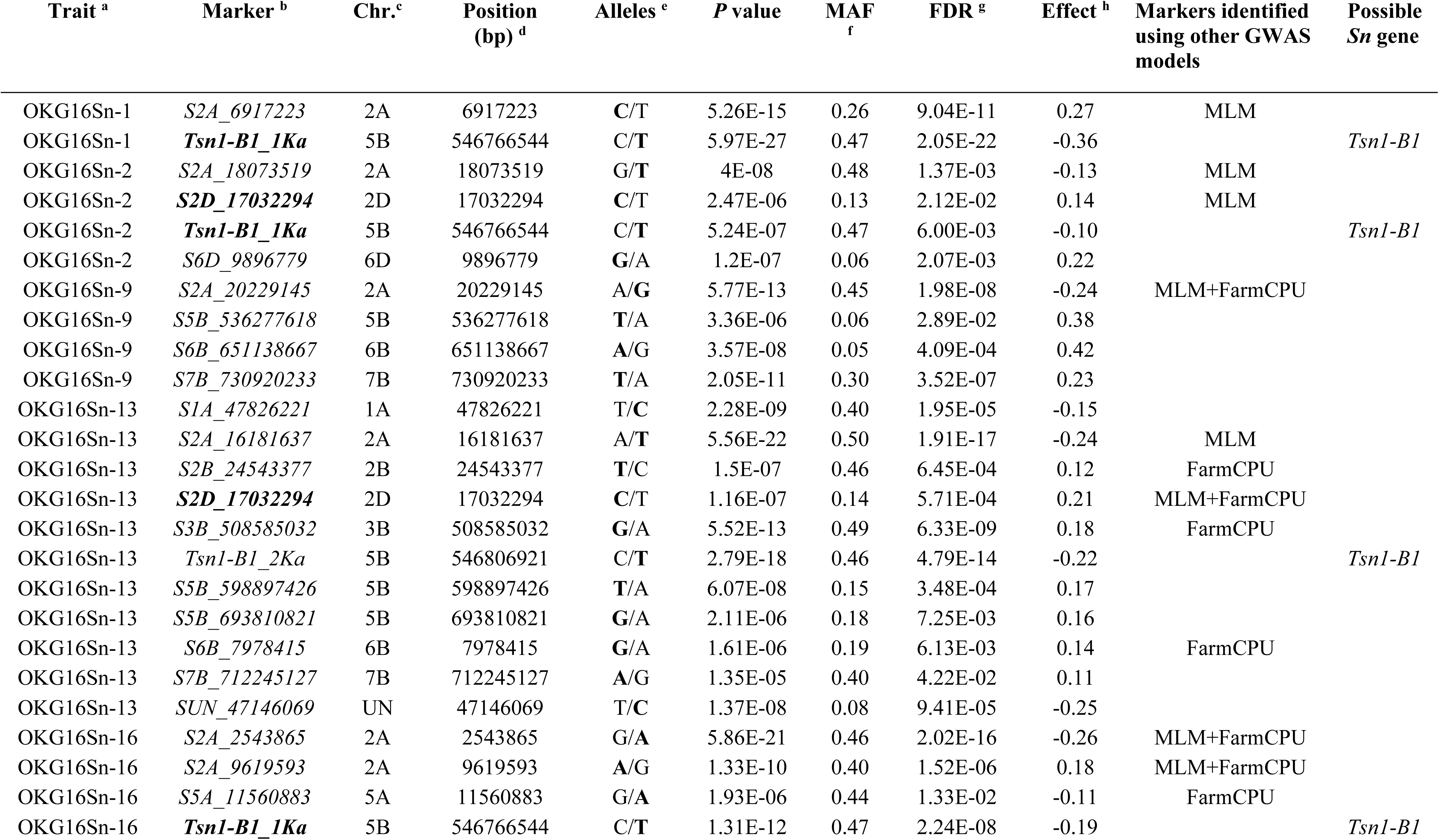
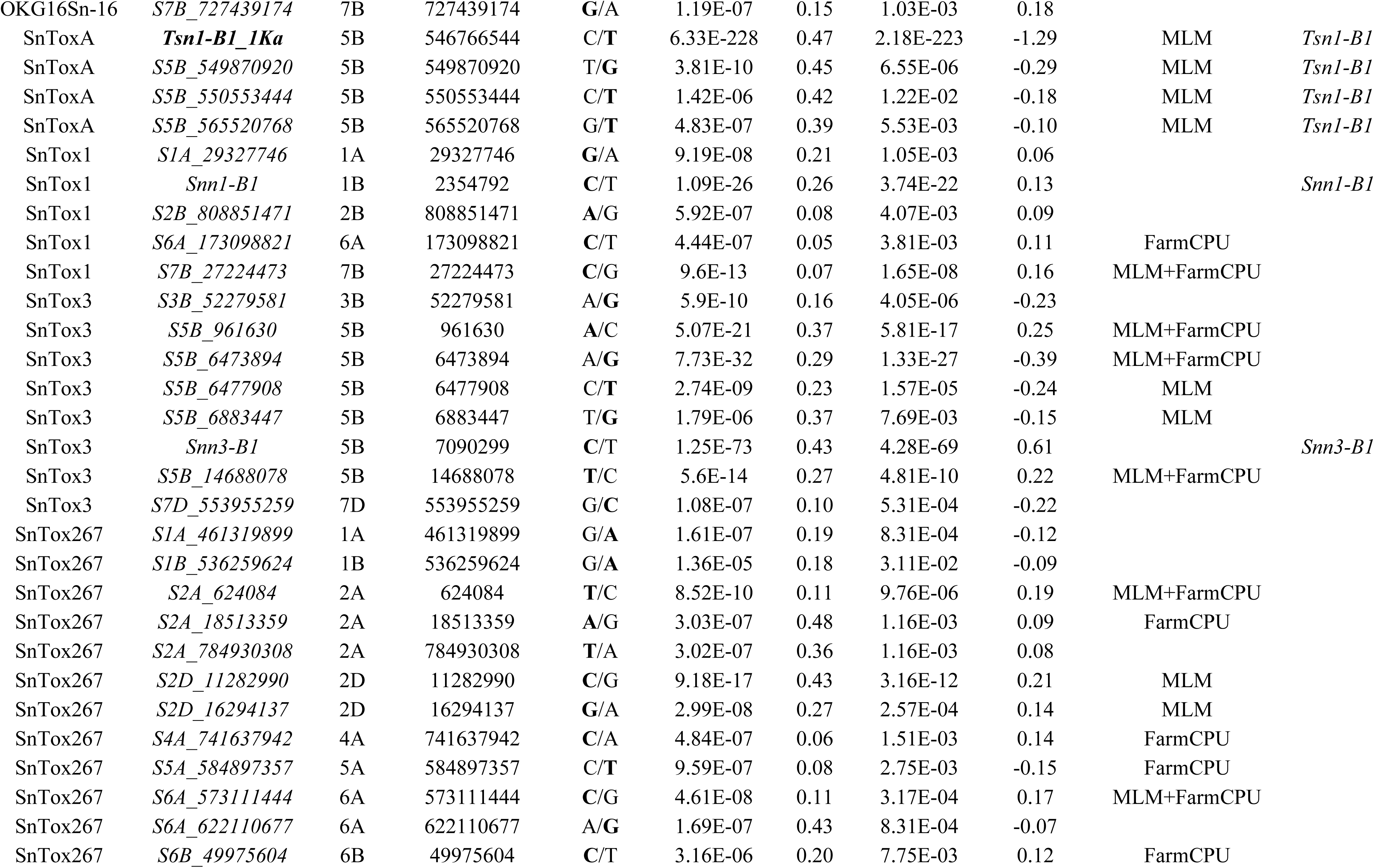
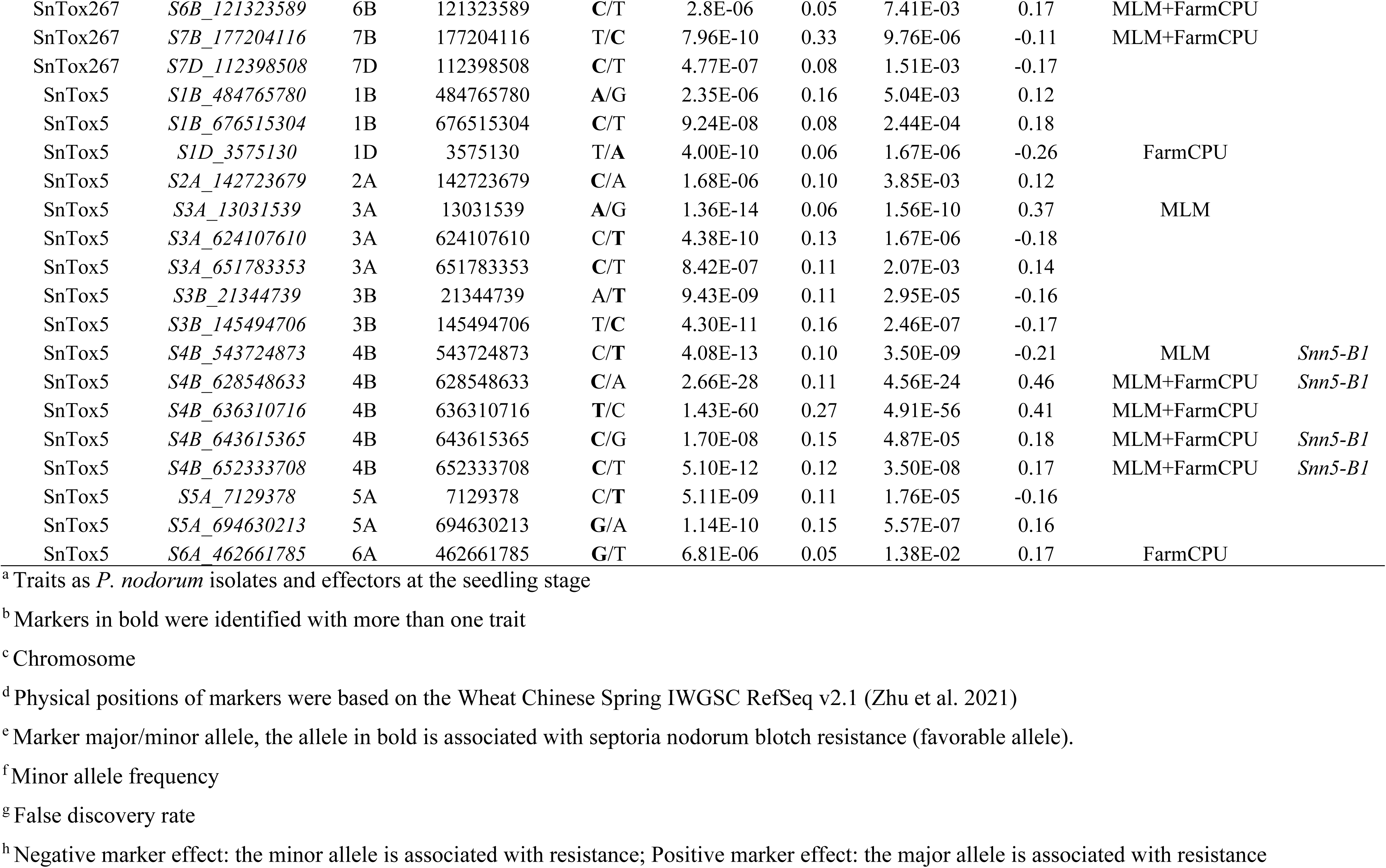
Summary of significant markers associated with septoria nodorum blotch response at the seedling stage using the BLINK model.

### Development and validation of KASP markers associated with response to septoria nodorum blotch

As no diagnostic marker for *Snn5-B1* is currently available, we targeted the region of *Snn5-B1* gene on chromosome arm 4BL by selecting the most significant SNP markers from the three tested GWAS models for KASP development (*S4B_636310716*, *S4B_628548633*, *S4B_652333708*, and *S4B_643615365*). Of these four SNPs, we successfully developed a KASP marker for *S4B_643615365*, here designated as *KASP_ S4B_643615365*. The primer sequences for the *KASP_ S4B_643615365* are provided in Table 5. *KASP_ S4B_643615365* was validated on 40 genotypes from this GWAS panel, with 20 resistant genotypes carrying the allele (C) and 20 susceptible genotypes carrying the allele (G) (Figure 5a). Genotype calls for the *KASP_ S4B_643615365* agreed with that of the GBS-SNP marker *S4B_643615365* for all the 40 tested genotypes.

**Fig. 5.**
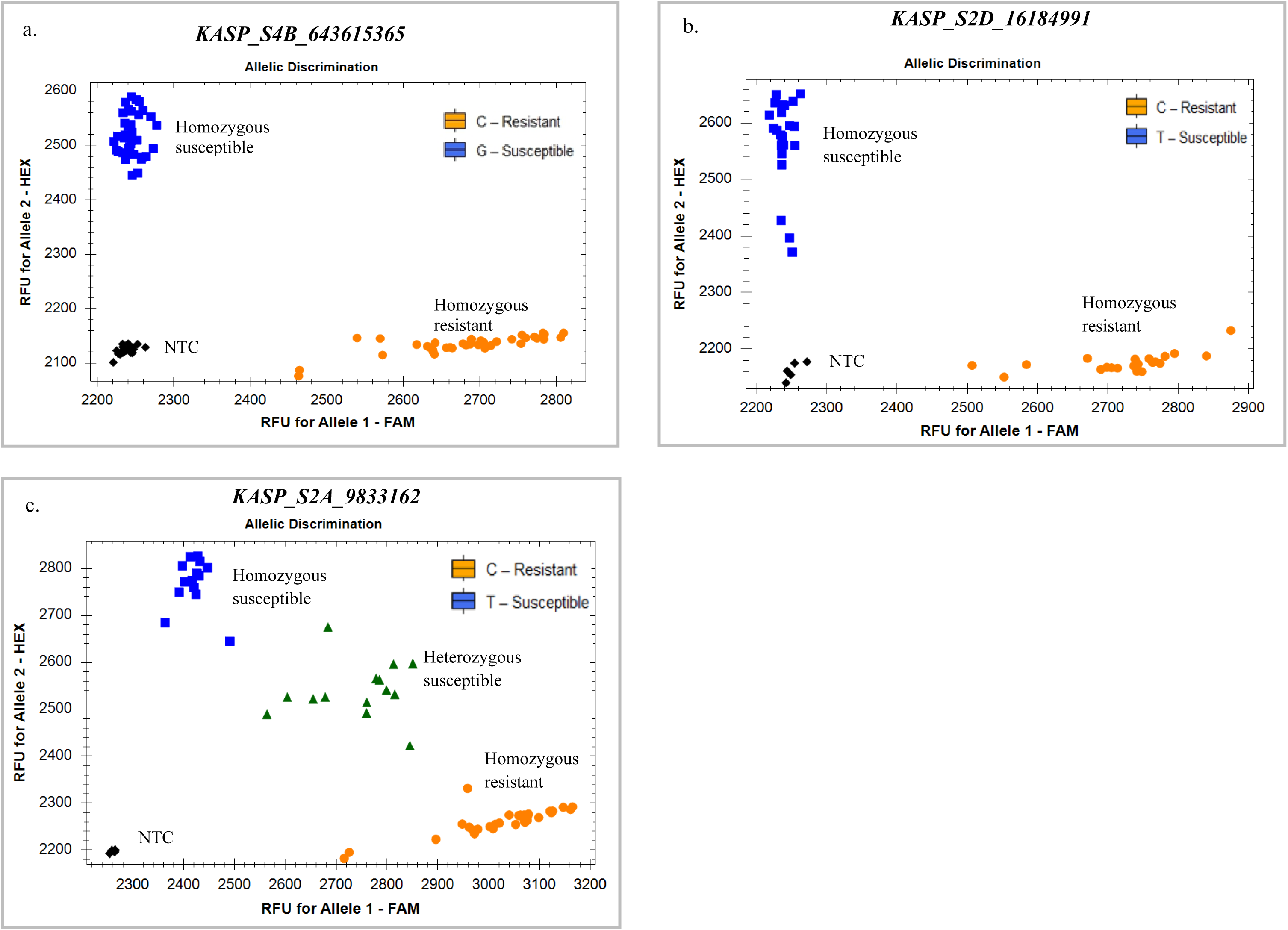
Genotyping scatter plots for the kompetitive allele-specific PCR (KASP) markers, *KASP_S4B_643615365*, *KASP_ S2D_16184991,* and *KASP_S2A_9833162* linked to *Snn5-B1*, *Snn2*, and *Qsnb.osu-2AS*, respectively.

**Table 5.**
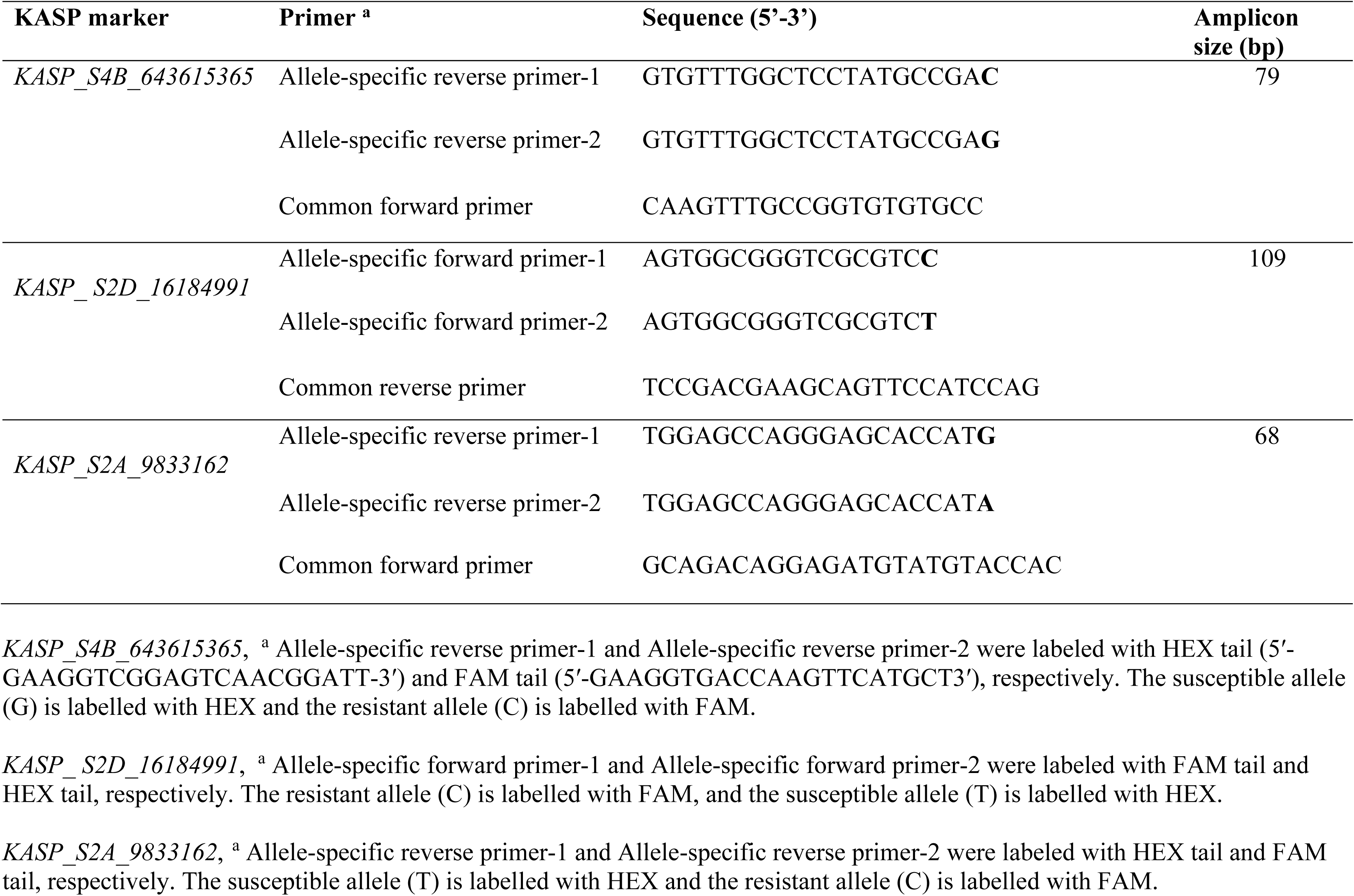
Kompetitive allele-specific PCR (KASP) marker developed from the most significant GBS SNP marker linked to *Snn5-B1*, *Snn2*, and *Qsnb.osu-2AS*.

As no diagnostic marker for *Snn2* is currently available, we targeted the locus of the *Snn2* gene on chromosome arm 2DS by selecting the most significant SNP markers from the three tested GWAS models for KASP development (*S2D_16294137*, *S2D_16383299*, *S2D_16184991*, and *S2D_16963560*). Of these four SNPs, we successfully developed a KASP marker for *S2D_16184991*, here designated as *KASP_ S2D_16184991*. The primer sequences for the *KASP_ S2D_16184991* were provided in Table 5. *KASP_ S2D_16184991* was validated on 22 genotypes from the OSU GWAS panel, with 11 resistant genotypes carrying the allele (C) and 11 susceptible genotypes carrying the allele (T) (Figure 5b). Genotype calls for the *KASP_ S2D_16184991* agreed with those of the GBS-SNP marker *S2D_16184991* for all 22 tested genotypes.

The novel locus on chromosome arm 2AS, designated here as *Qsnb.osu-2AS*, was found to be associated with response to all five *P. nodorum* isolates tested in this study. To develop KASP markers, we targeted the most significant SNPs on chromosome 2AS from the three tested GWAS models for KASP development (*S2A_9619593*, *S2A_2543865*, *S2A_9833162*, and *S2A_17970052*). Of these four SNPs, we successfully developed a KASP marker for *S2A_9833162*, here designated as *KASP_ S2A_9833162*. The primer sequences for the *KASP_ S2A_9833162* were provided in Table 5. *KASP_ S2A_9833162* was validated on 60 genotypes from the OSU GWAS panel, with 30 resistant genotypes carrying the allele (C) and 30 susceptible genotypes carrying the allele (T) (Figure 5c). Genotype calls for the *KASP_ S2A_9833162* agreed with that of the GBS-SNP marker *S2A_9833162* for all 30 tested resistant genotypes. The susceptible genotypes showed a mixture of homozygous susceptible alleles or heterozygous calls.

## Discussion

The increased incidence of septoria nodorum blotch in HWW production regions in the Great Plains (Aoun and Carver 2024) has made breeding for SNB resistance a priority. This is the first comprehensive study of SNB susceptibility/resistance in contemporary HWW, integrating multiple approaches: evaluating host reactions to *P. nodorum* isolates and NEs, identifying characterized SNB sensitivity genes using diagnostic markers, and validating known sensitivity genes while uncovering novel resistance/susceptibility loci through genome-wide association studies (GWAS). Seedling evaluations of 619 HWW breeding lines and cultivars to *P. nodorum* isolates showed that up to 67% of the genotypes were susceptible. Richards et al. (2019) found that *P. nodorum* isolates collected from Oklahoma were clustered separately from all other isolates collected from other U.S. states. Furthermore, all Oklahoma isolates in this survey carried *SnToxA*, which interacts with *Tsn1* to cause susceptibility. In this study, we found that 54% of this HWW panel were sensitive to SnToxA, thus the frequency of *Tsn1* should be reduced in HWW to enhance SNB resistance. *Tsn1*, which encodes a protein with integrated serine/threonine protein kinase (PK), nucleotide binding, and leucine-rich repeat (NLR) domains (Faris et al. 2010), also causes susceptibility to other fungal diseases, including tan spot (*Pyrenophora tritici-repentis*) and spot blotch (*Bipolaris sorokiniana*), making it the first sensitivity gene to target with marker-assisted elimination. Previous studies showed *Tsn1* is present in 32% - 76% of spring and winter wheat panels (Waters et al. 2011; Tan et al. 2014; Bertucci et al. 2014; Ruud et al. 2018; Hafez et al. 2020; AlTameemi et al. 2021; Peters Haugrud et al. 2023; Szabo-Hever et al. 2025b). The high frequency of *Tsn1* in different wheat classes is likely due to its genetic linkage with other genes that confer desirable traits in wheat, or this gene may have a function other than conferring sensitivity to SnToxA (Richards et al. 2019; Cowger et al. 2020; Hafez et al. 2020; AlTameemi et al. 2021; Szabo-Hever et al. 2025a). Faris et al. (2010b) demonstrated that the *Tsn1* gene originated from a gene fusion event that might be functioning in the diploid B-genome progenitor of polyploid wheat. Consequently, *Tsn1* was perhaps transferred to hexaploid wheat from tetraploid wheat by either amphiploidization or secondary hybridization events.

Sensitivity to SnTox3 was present in 37% of this HWW panel, making *Snn3* another target gene to eliminate in HWW breeding programs. The frequency of SnTox3 sensitivity was lower compared to that (52%) reported in a hard red spring wheat panel and higher compared to that (22%) observed in a global durum wheat panel (Szabo-Hever et al. 2025b). *Tsn1*-*SnToxA* and *Snn3*-*SnTox3* interactions were found to play an important role in SNB sensitivity in other spring and winter wheat panels (Adhikari et al. 2011; Gurung et al. 2014; Liu et al. 2015; Phan et al. 2018; Ruud et al. 2019; Halder et al. 2019; AlTameemi et al. 2021; Peters Haugrud et al. 2023).

The frequency of SnTox1 sensitivity was low (2%) in this HWW panel, which is lower than that observed in panels of hard red spring wheat (28%; Szabo-Hever et al. 2025b), winter wheat (37%; Peters Haugrud et al. 2023), and durum wheat (65%; Szabo-Hever et al. 2025b). The low frequency of SnTox1 could be due to the absence of a functional *Snn1-B1* allele, which was evident based on the false positive rate of 25% of the diagnostic marker available for this allele. *Snn1* could have gone through mutation or deletion, resulting in the loss of its sensitivity; thus, the available *Snn1-B1* diagnostic marker could not detect these diverse structural variations (Seneviratne et al. 2024a). Shi et al. (2016) showed that a homoeoallele of *Snn1* is present on chromosome 1D of hexaploid wheat, which could compensate for the loss of *Snn1* on 1B. The lack of significant associations on chromosome 1D for response to SnTox1, could be due to the low allele frequency of the D-genome copy of *Snn1*.

Sensitivity to SnTox267 and SnTox5 in this HWW panel was 13% and 15%, respectively. These frequencies were lower compared to those reported in a winter wheat panel, which were 39% and 42% sensitivity to SnTox267 and SnTox5, respectively (Peters Haugrud et al. 2023). Szabo-Hever et al. (2025b) reported that sensitivity to SnTox267and SnTox5 in a hard red spring wheat panel was 54% and 12%, respectively. Our GWAS in response to SnTox267 identified significant associations linked to *Snn2* (on chromosome 2DS; Friesen et al. 2007), but not to *Snn6* (on chromosome 6AL; Gao et al. 2015) and *Snn7* (on chromosome 2DL; Shi et al. 2015). These results agreed with those of Peters Haugrud et al. (2023), where no associations were found for *Snn6* and *Snn7* genes in a global winter wheat collection. In addition to *Snn2*, Szabo-Hever et al. (2025b) identified associations on chromosome 2DL for response to SnTox267 in hard red spring wheat, which may correspond to *Snn7*, but did not show any associations linked to *Snn6*. *Snn7* was identified and mapped in Chinese Spring’ (CS)–Timstein (CS-Tm) disomic chromosome substitution lines (Gao et al. 2015; Shi et al. 2015b). The sensitivity gene *Snn6* is not common in hexaploid wheat and was identified only in the synthetic wheat line W-7984 (Gao et al. 2015).

Our GWAS reported that sensitivity to SnTox5 was mainly attributed to the presence of *Snn5-B1* on chromosome 4BL. As no diagnostic markers are currently available to eliminate *Snn5*, a KASP marker (*KASP_ S4B_643615365*) was developed from a significant SNP (*S4B_643615365*) on chromosome 4BL, which has an accuracy of 89% in predicting sensitivity/insensitivity to SnTox5 (false positive rate = 7.3% and false negative rate= 36%). The GWAS BLINK model also identified other loci on chromosome 1B, 1D, 2A, 3A, 3B, 5A, and 6A to be associated with response to SnTox5. Additional loci (other than *Snn5*) on chromosomes 5A and 5B were also found to be associated with response to SnTox5 in winter wheat (Peters Haugrud et al. 2023). Further work is needed to determine the interaction between SnTox5 and any of these GWAS identified loci.

Although isolate OKG16Sn-9 did not produce the NEs SnTox1 or SnTox267, it produced the highest disease scores on susceptible lines, reaching a severity score of 5 (a level not observed with any of the other isolates). Phan et al. (2016) reported that the *Snn1*−SnTox1 interaction is epistatic with *Snn3*−SnTox3 interaction, and higher expression of SnTox3 in *P. nodorum* was observed in the absence of SnTox1. Peters Haugrud et al. (2019) also reported antagonistic epistasis between SnTox1 and SnToxA, where *SnTox1* gene expression levels in isolate Sn2000 was lower when SnToxA was produced. This may explain the lack of significant GWAS associations in the locus of *Snn1* on chromosome 1BS for responses to all five *P. nodorum* isolates (SnToxA present in all isolates) in this study. Our findings also suggest the involvement of previously unidentified necrotrophic effectors (NEs) in *P. nodorum*, as well as the presence of uncharacterized sensitivity/resistance genes in wheat. This was evident based on the GWAS, where unknown loci were identified for responses to the five isolates and NEs (except ToxA). Of which, a locus on chromosome 2AS, *Qsnb.osu-2AS*, was found to be associated with responses to all five isolates and had higher effects compared to *Tsn1* gene for responses to all isolates except OKG16Sn-1 and OKG16Sn-9. The HWW variety ‘Green Hammer’, which is currently the second most grown variety in Oklahoma (USDA NASS 2025) was insensitive to all five NEs and did not carry any of the corresponding sensitivity genes but was susceptible to isolates OKG16Sn1-1 and OKG16Sn-2. This suggests that other unreported sensitivity genes targeted by known or novel NEs could be present in Green Hammer and other NEs could be identified in OKG16Sn1-1 and OKG16Sn-2. In addition to the known sensitivity genes in wheat, our GWAS identified novel loci to be associated with responses to *P. nodorum* isolates and NEs. This suggests that other NEs and resistance/sensitivity SNB genes could be identified in the *P. nodorum*-wheat system. As *Tsn1-B1* and *Snn3-B1/B2* are present in high frequencies in contemporary HWW, their elimination should be prioritized in HWW breeding programs using available and accurate markers. We developed KASP markers linked to *Snn5-B1*, *Snn2*, and *Qsnb.osu-2AS*.

### Statements and declarations

This project was funded by the U.S. Department of Agriculture National Institute of Food and Agriculture Grant # 2024-67013-42587. The mention of trade names or commercial products in this publication is solely to provide specific information and does not imply recommendation or endorsement by the United States Department of Agriculture. The USDA is an equal opportunity provider and employer.

## Data availability statement

All data generated analyzed during this study are included in this published article and its supplementary information files submitted with this manuscript. Supplementary Tables (Table S1–S8) and Supplementary Figures (Figure S1 – S9) are available at https://doi.org/10.6084/m9.figshare.30531503.v4

