## Supplementary Figures for "Identification of septoria nodorum blotch susceptibility genes in hard winter wheat"

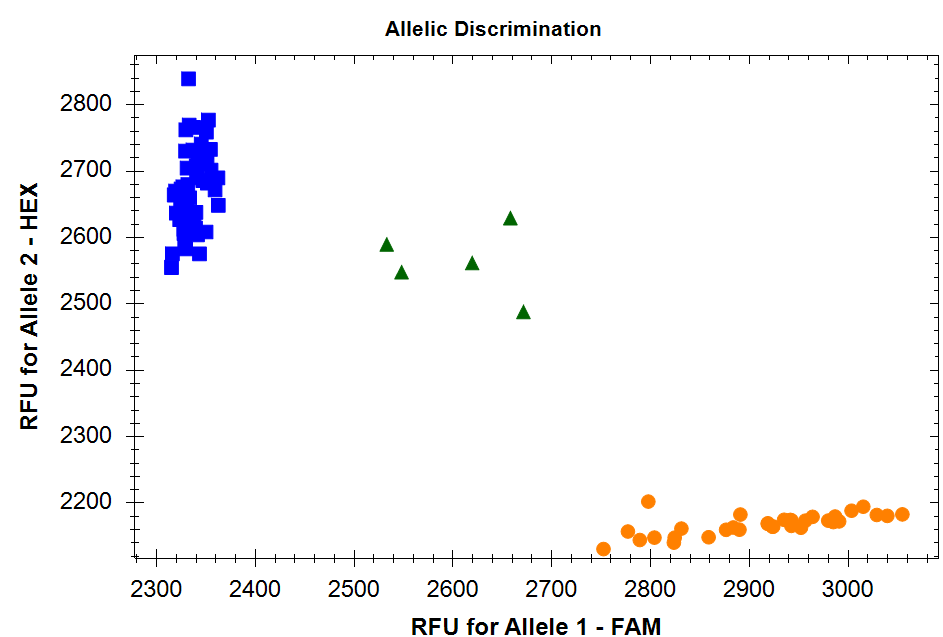

Tsn1-B1_1Ka

*Tsn1-B1+*

*Tsn1-B1-*

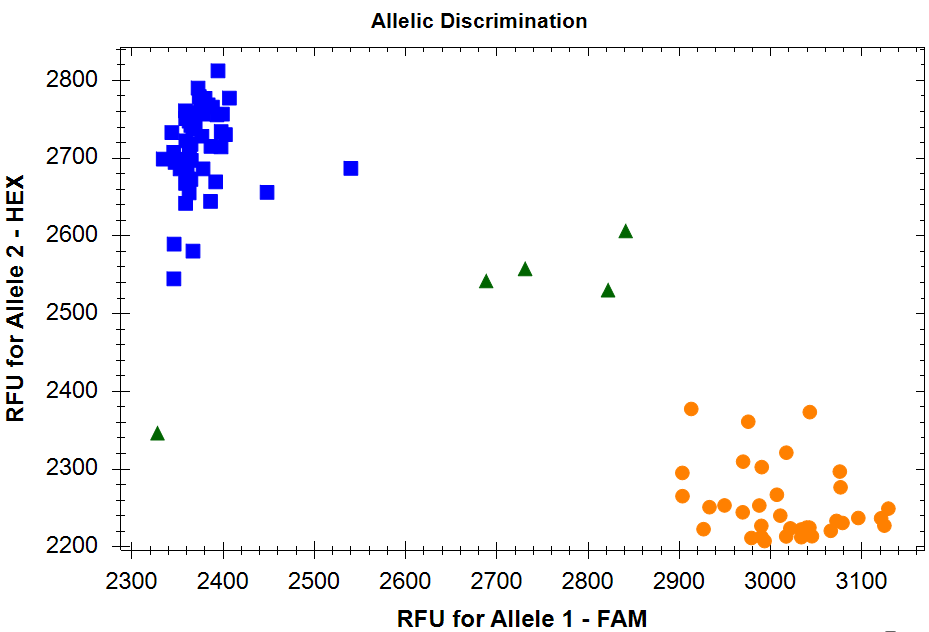

Tsn1-B1_2Ka

*Tsn1-B1+*

*Tsn1-B1-*

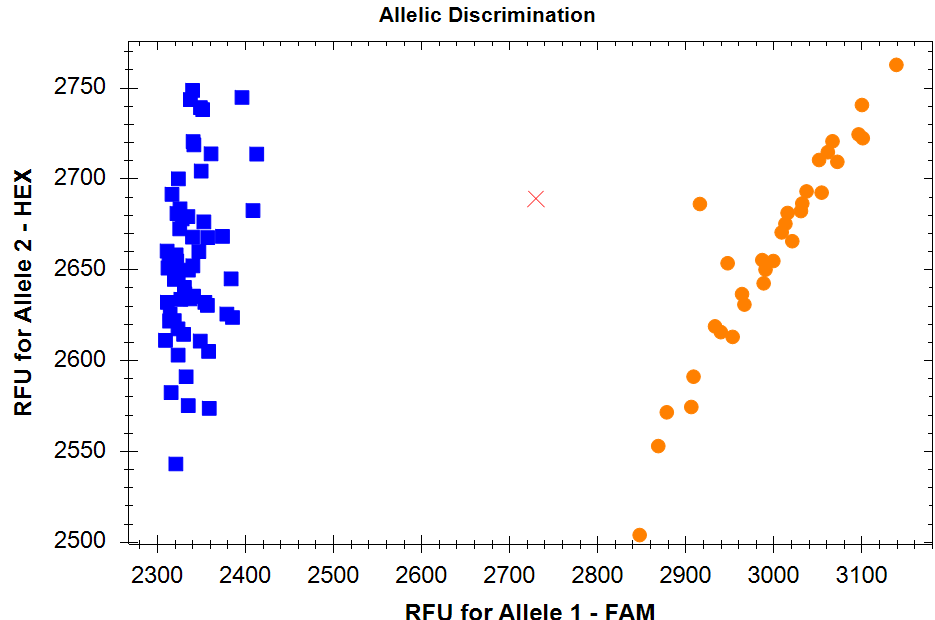

*Snn1*

*Snn1+*

*Snn1-*

Snn1_null

*Snn1-B1-*

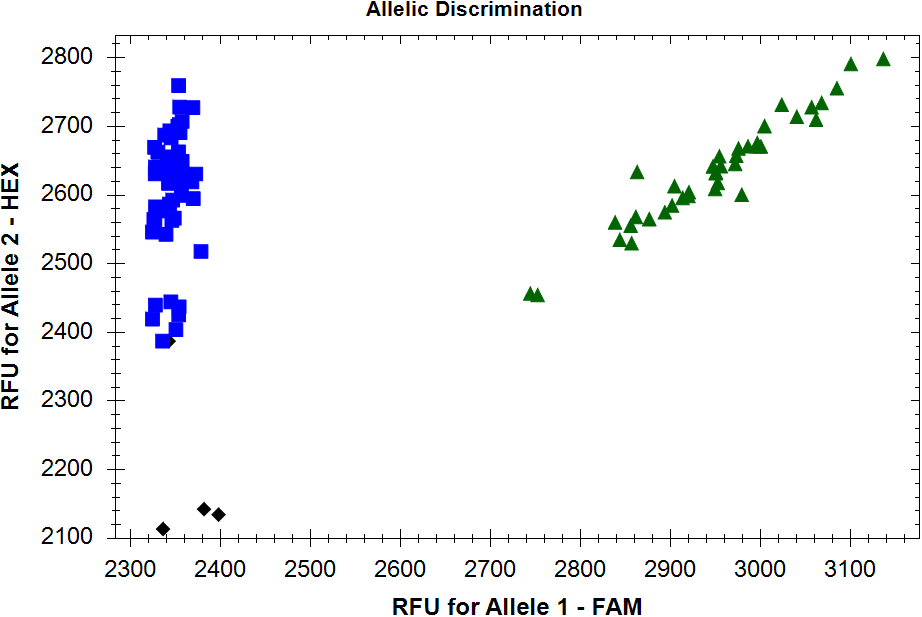

Snn3-B2

*Snn3-B2-*

*Snn3-B2+*

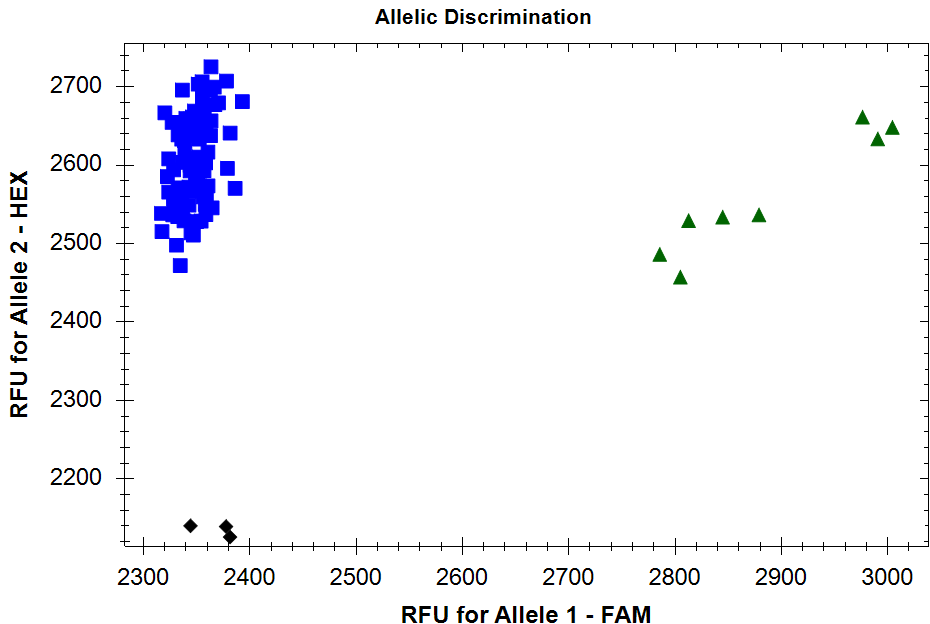

Snn3-B1

*Snn3-B1+*

*Snn3-B1-*

a)

b)

c)

d)

*Snn1-B1+*

**Supplementary Fig. S1**. Scatter plots showing genotyping results of the hard winter genotypes using available kompetitive allele-specific PCR (KASP) markers for the sensitivity genes *Tsn1-B1*, *Snn1-B1*, *Snn3-B1*, and *Snn3-B2*. (a) *Tsn1_1Ka* and *Tsn1_2Ka* both markers identify the presence (+) or absence (-) of *Tsn1-B1*. (b) *Snn1_null* marker identifies the presence/absence of *Snn1-B1* gene. (c) *Snn3-B1* marker identifies the presence/absence of *Snn3-B1* gene. (d) *Snn3-B2* marker identifies the presence/absence of *Snn3-B2* gene.

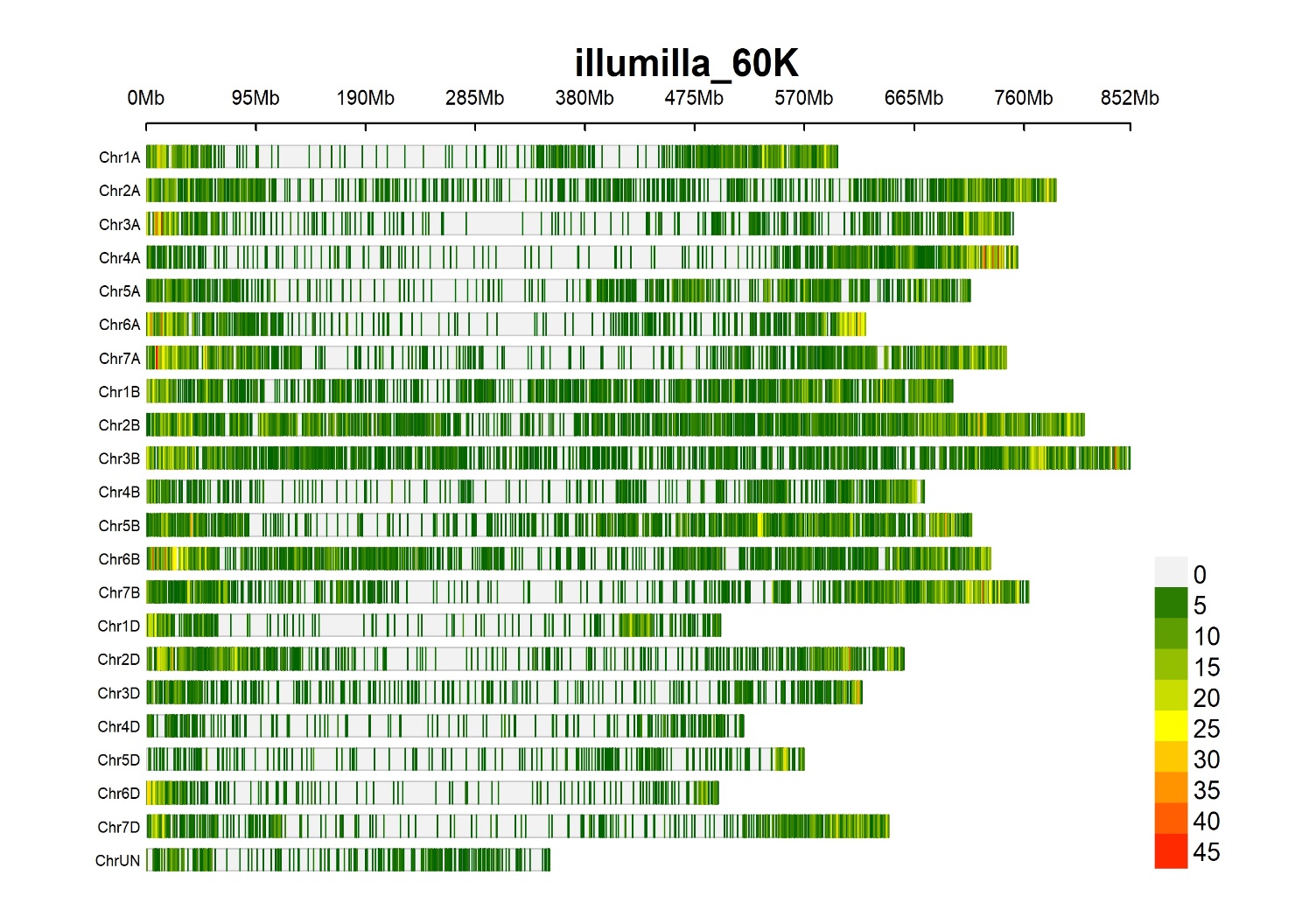

**The SNP markers density within 1Mb window on wheat chromosomes**

**Supplementary Fig. S2**. Density of SNP markers (n = 34,357 SNPs) per 1Mb window on wheat chromosomes in a set of 619 hard winter wheat genotypes.

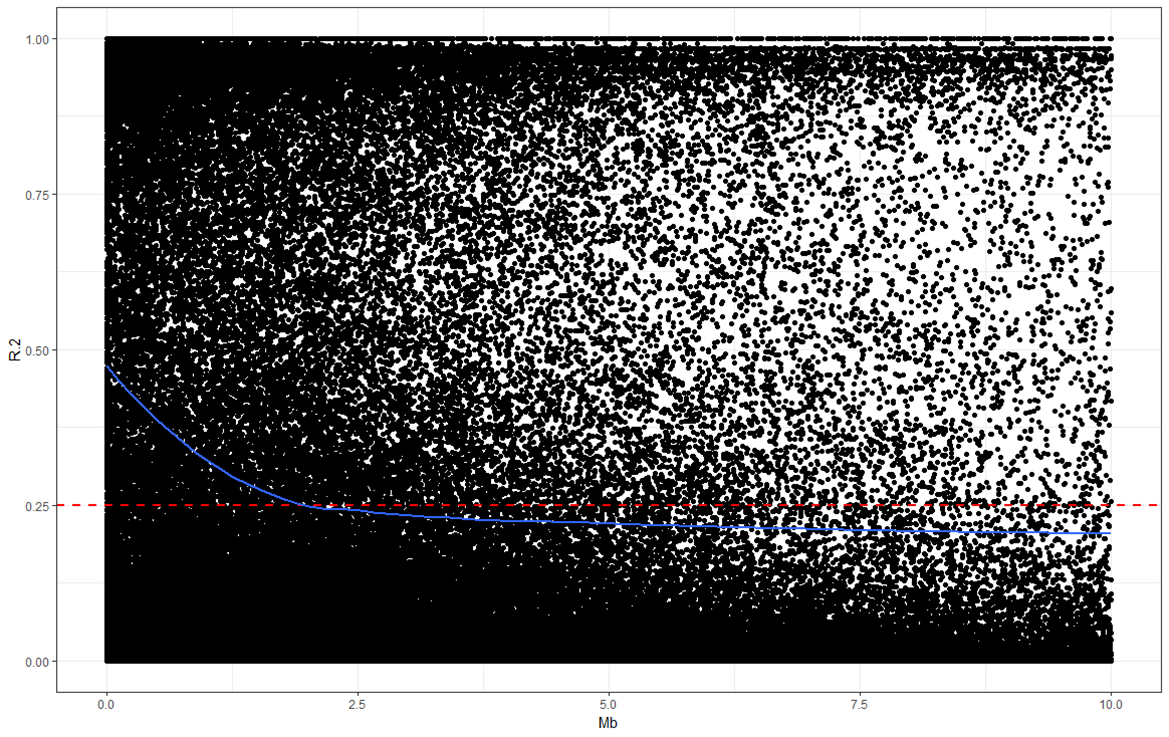

**Supplementary Fig. S3.** Scatter plot showing linkage disequilibrium (LD) decay across the whole genome. The LD estimates (*r*^2^) for pairs of SNPs were plotted against the corresponding physical positions in million base pair (Mb) based on the Chinese Spring wheat reference genome IWGSC_RefSeqv2.1 (Zhu et al. 2021).

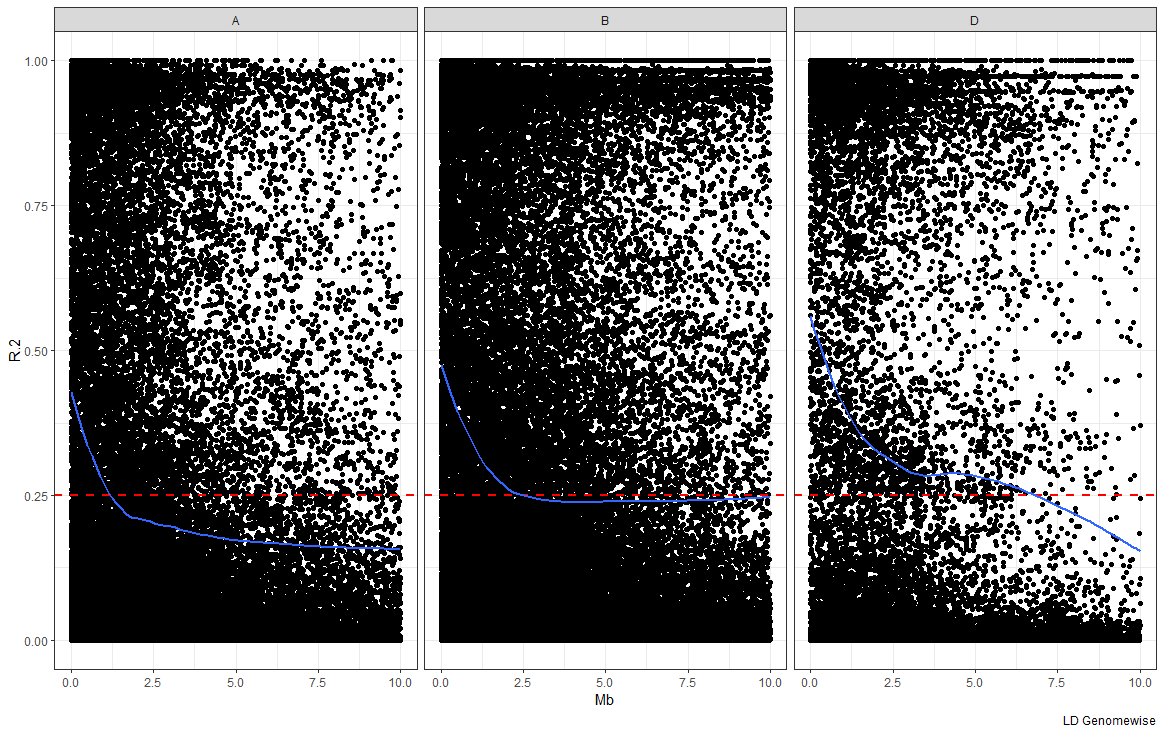

**Supplementary Fig. S4**. Scatter plot showing linkage disequilibrium (LD) decay in genomes A, B, and D. The LD estimates (*r*^2^) for pairs of SNPs were plotted against the corresponding physical positions in million base pair (Mb) based on the Chinese Spring wheat reference genome IWGSC_RefSeqv2.1 (Zhu et al. 2021).

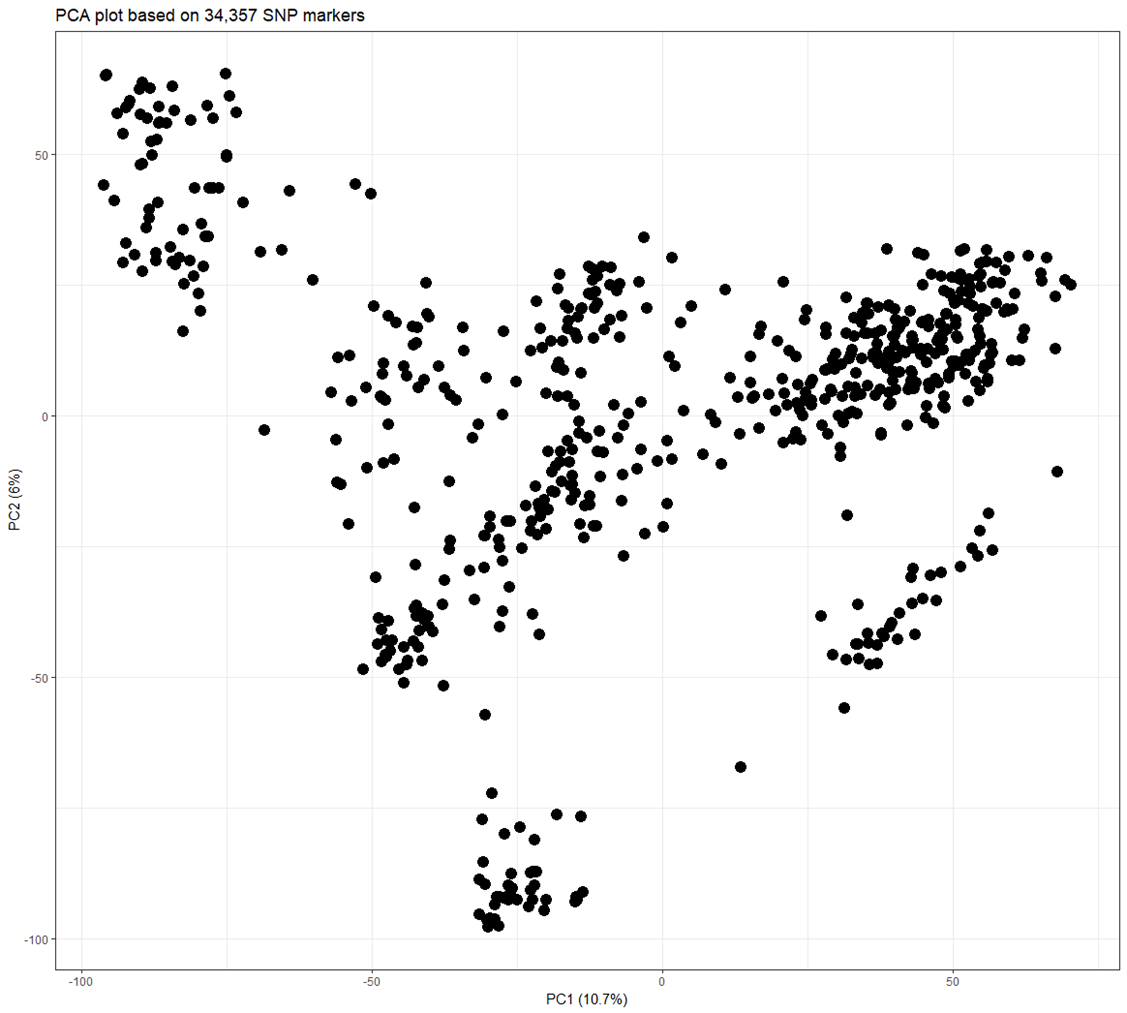

**Supplementary Fig. S5.** Principal component (PC) analysis obtained from 34,357 single nucleotide polymorphisms (SNPs) in 619 hard winter wheat genotypes. The first two PCs, PC1 and PC2, explained 10.7% and 6.0% of the variation, respectively.

**OKG16Sn-1**

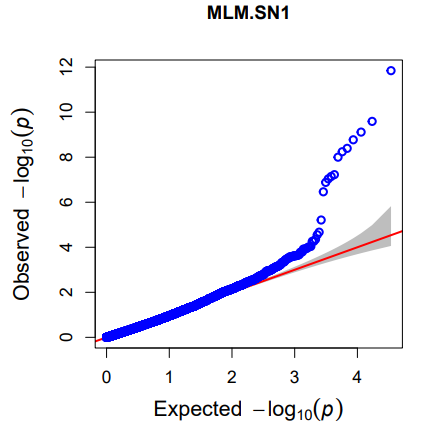

MLM (K+PC0)

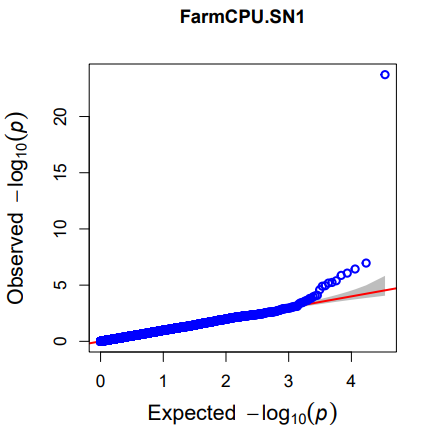

FarmCPU (K+PC2)

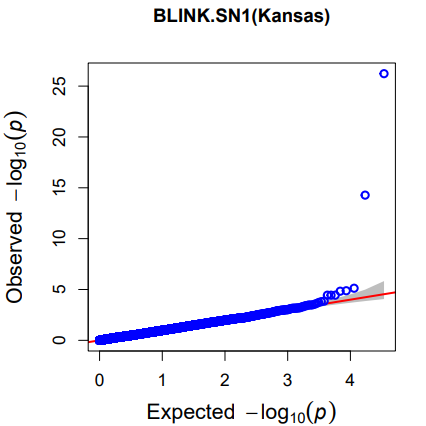

BLINK(K+PC0)

**OKG16Sn-2**

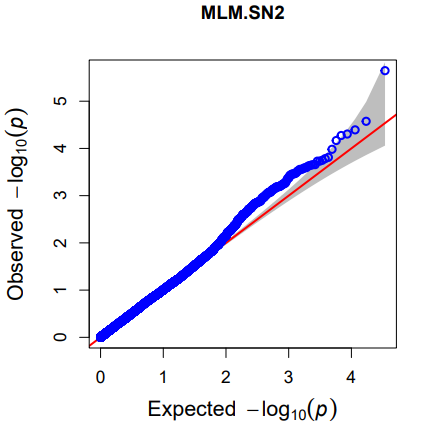

MLM (K+PC4)

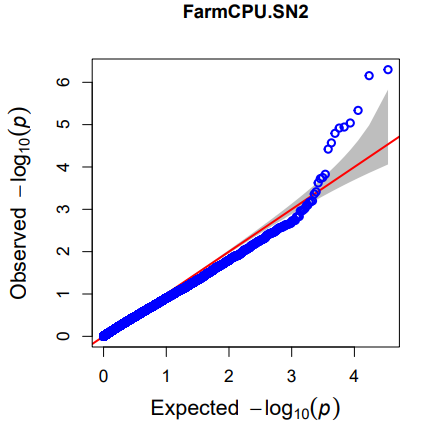

FarmCPU (K+PC2)

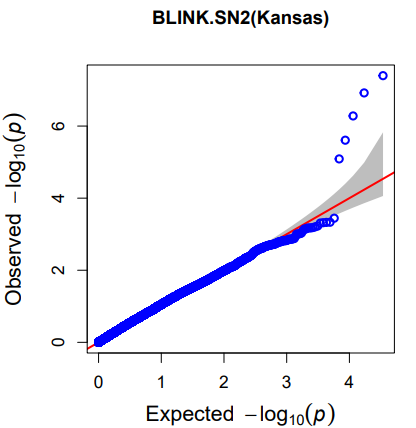

BLINK(K+PC4)

**OKG16Sn-9**

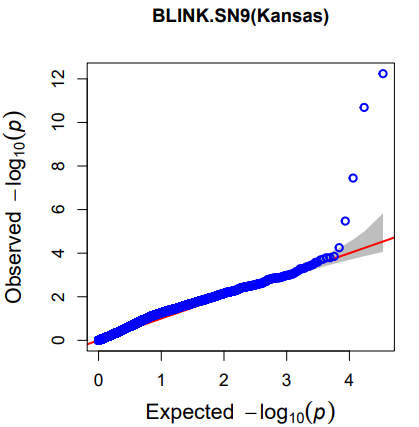

BLINK (K+PC0)

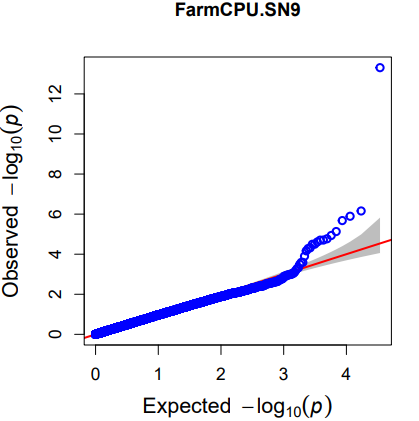

FarmCPU (K+PC2)

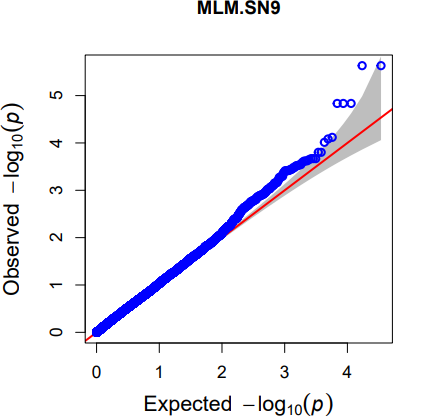

MLM (K+PC4)

**OKG16Sn-13**

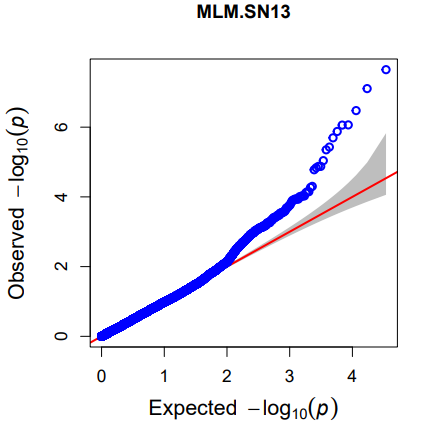

MLM (K+PC0)

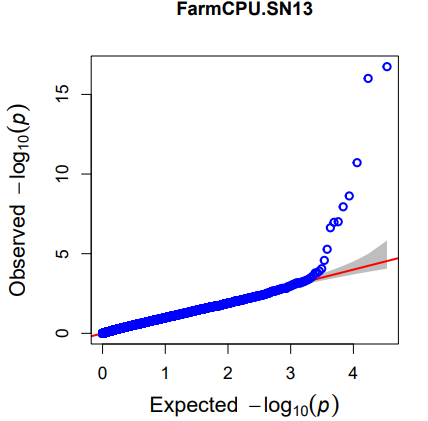

FarmCPU (K+PC2)

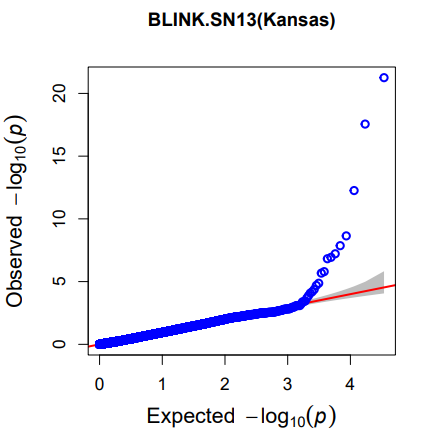

BLINK (K+PC0)

**OKG16Sn-16**

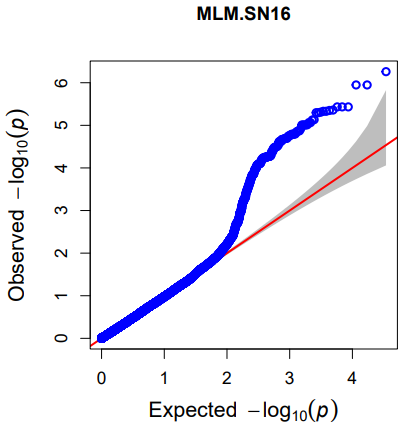

MLM (K+PC0)

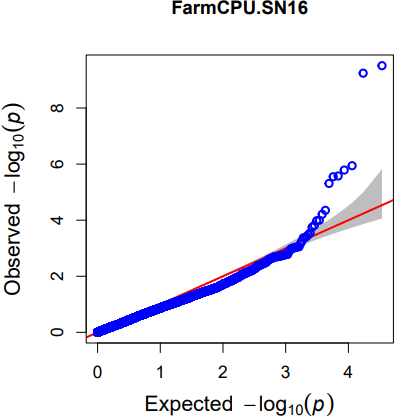

FarmCPU (K+PC0)

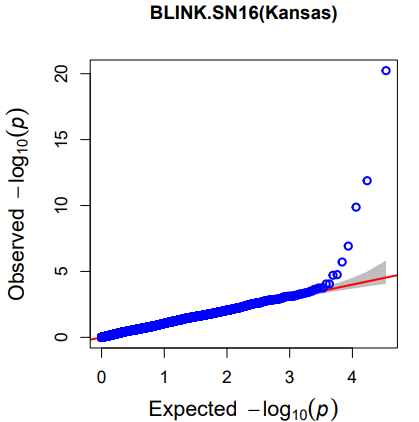

BLINK (K+PC0)

**Supplementary Fig. S6**. Quantile-quantile (Q-Q) plots comparing the expected -log10 (*P*) values versus the observed -log10 (*P*) values of the selected association mapping models for responses to five *P. nodorum* isolates. Within MLM, FarmCPU, and BLINK models, family relatedness (K matrix) and up to four principal components (PCs) in population structure (Q matrix) were tested.

**SnToxA**

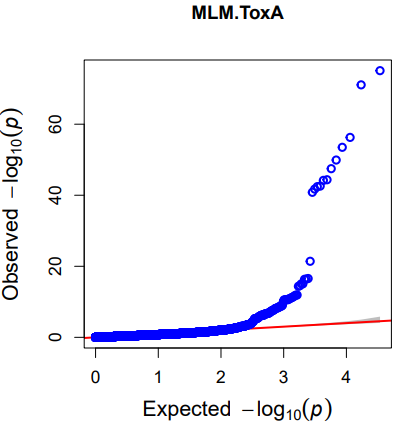

MLM (K+PC0)

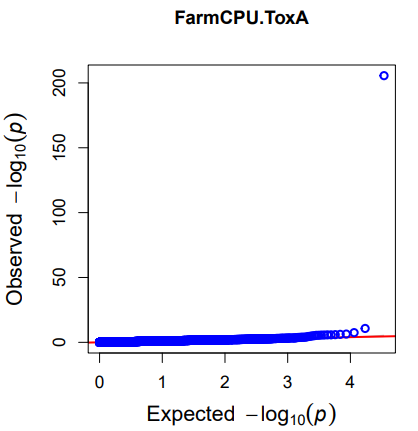

FarmCPU (K+PC0)

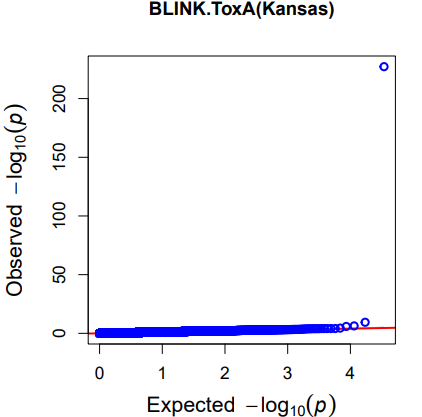

BLINK (K+PC0)

**SnTox1**

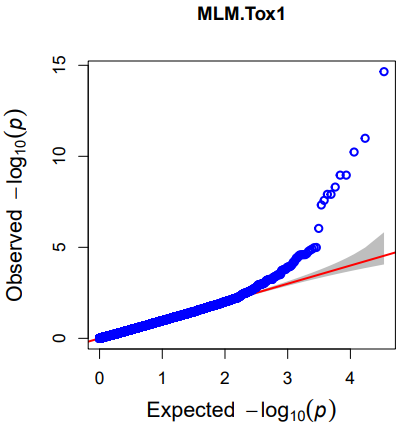

MLM (K+PC0)

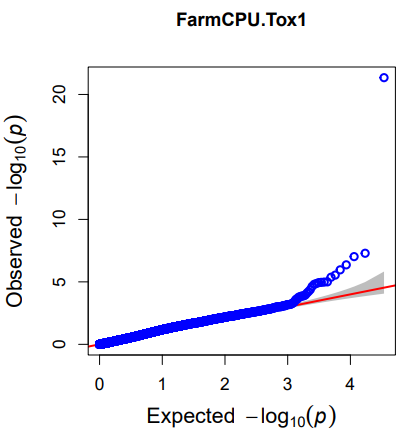

FarmCPU (K+PC0)

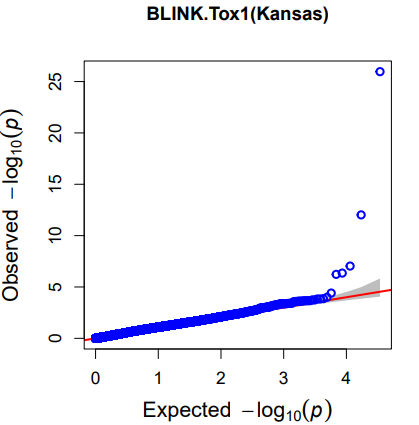

BLINK (K+PC0)

**SnTox3**

**SnTox267**

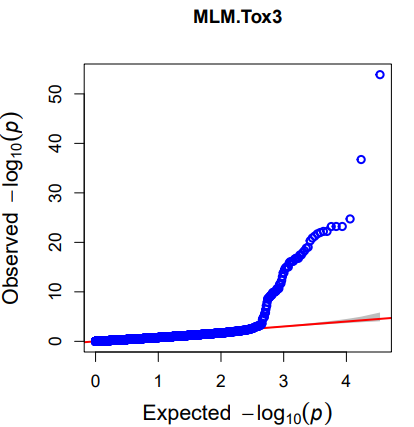

MLM (K+PC0)

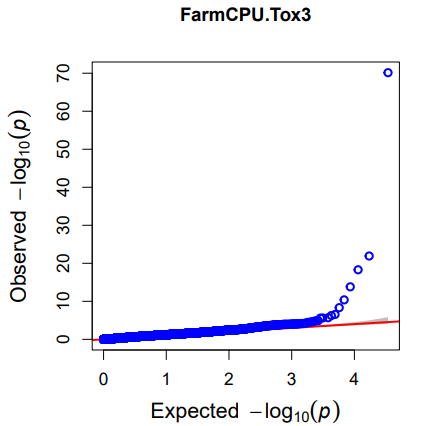

FarmCPU (K+PC0)

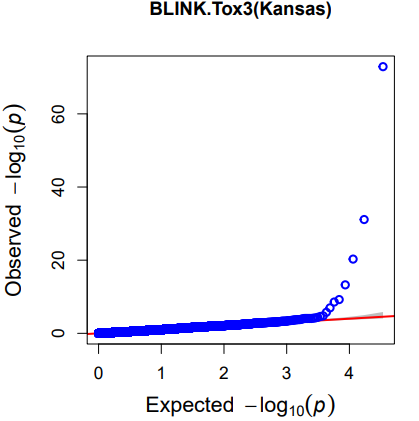

BLINK (K+PC2)

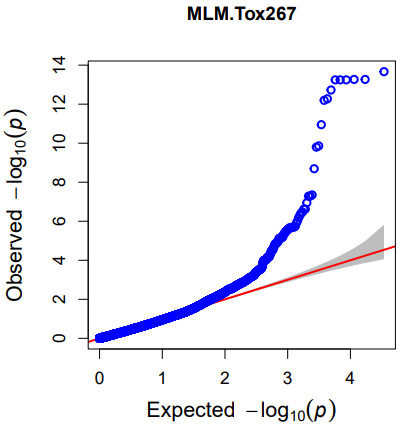

MLM (K+PC0)

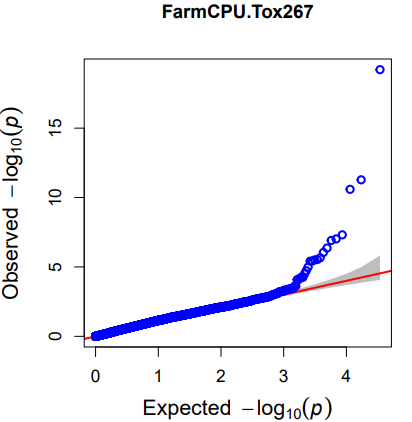

FarmCPU (K+PC2)

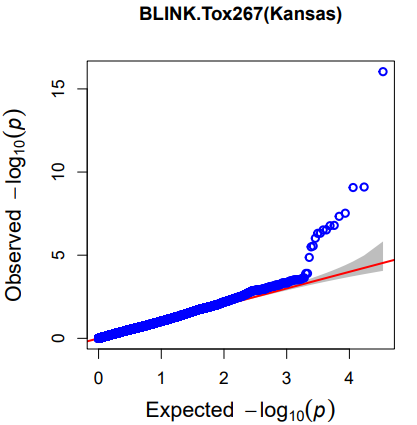

BLINK (K+PC2)

**SnTox5**

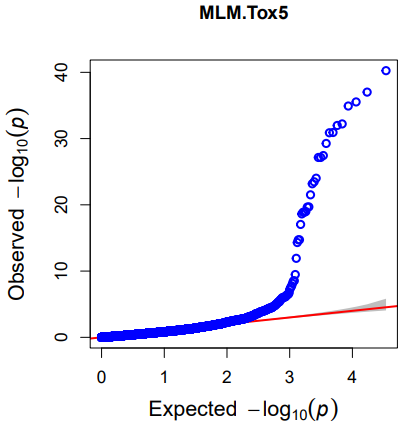

MLM (K+PC0)

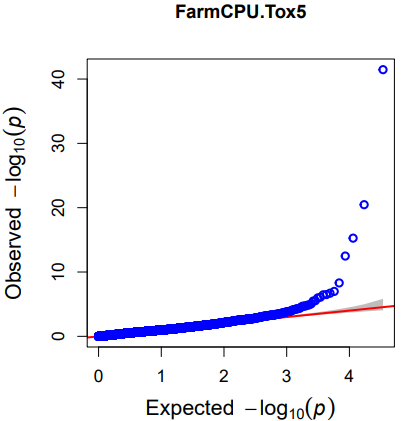

FarmCPU (K+PC0)

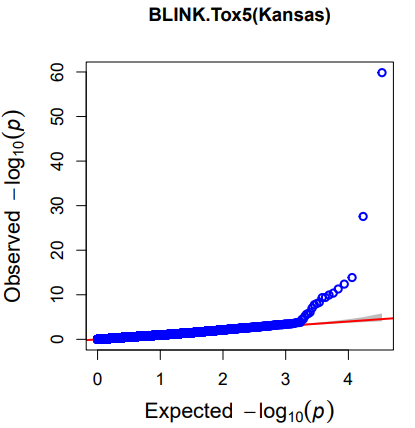

BLINK (K+PC0)

**Supplementary Fig. S7**. Quantile-quantile (Q-Q) plots comparing the expected -log10 (*P*) values versus the observed -log10 (*P*) values of the selected association mapping models for responses to five *P. nodorum* effectors. Within MLM, FarmCPU, and BLINK models, family relatedness (K matrix) and up to four principal components (PCs) in population structure (Q matrix) were tested.

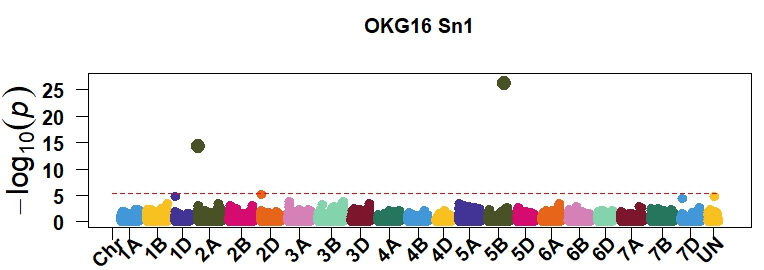

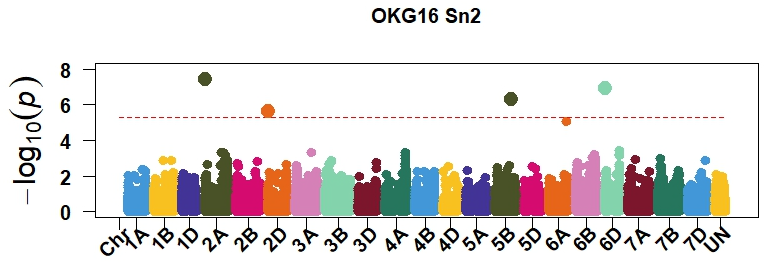

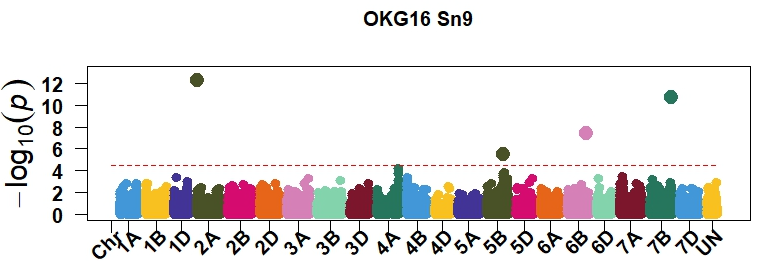

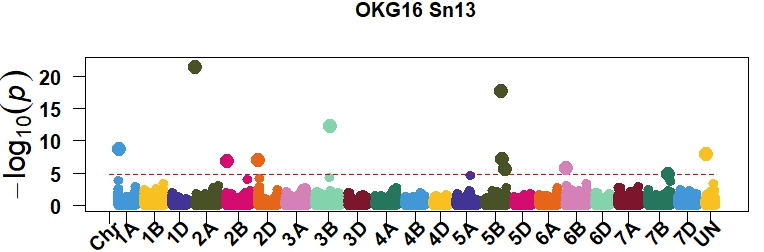

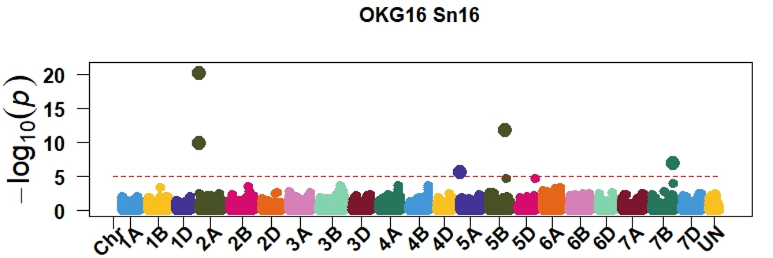

*Tsn1-B1*

OKG16Sn-1

OKG16Sn-13

OKG16Sn-2

OKG16Sn-16

OKG16Sn-9

*Tsn1-B1*

*Tsn1-B1*

*Tsn1-B1*

*Tsn1-B1*

**Supplementary Fig. S8**. Manhattan plots showing significant markers associated with responses to five *P. nodorum* isolates using the BLINK model. The horizontal red line indicates significance levels at a false discovery rate ≤ 0.05.

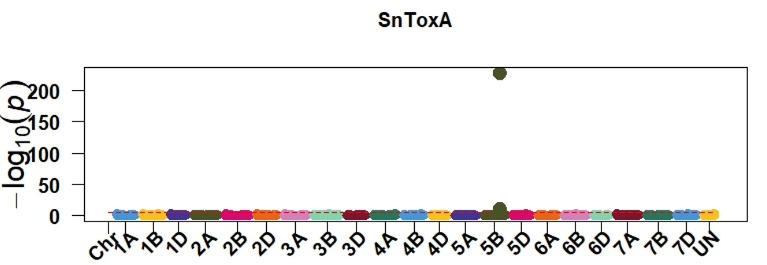

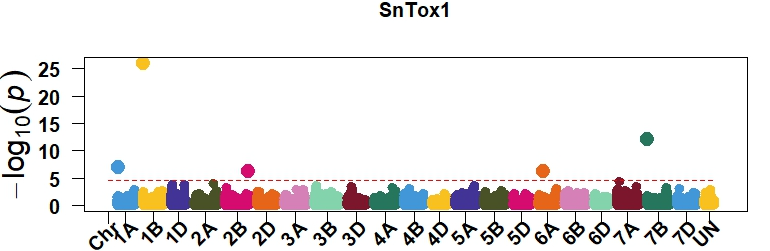

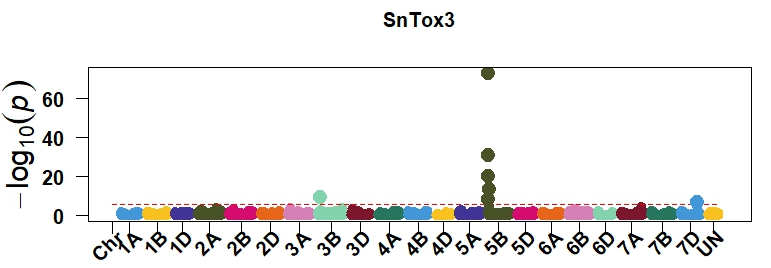

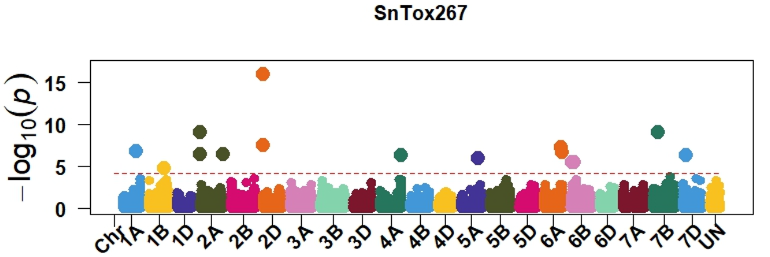

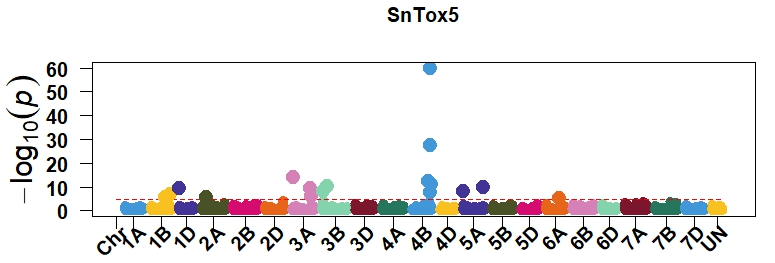

SnToxA

SnTox1

SnTox3

SnTox267

SnTox5

*Tsn1-B1*

*Snn1-B1*

*Snn3-B1*

*Snn2*

*Snn5-B1*

**Supplementary Fig. S9.** Manhattan plots showing significant markers associated with five *P. nodorum* effectors using the BLINK model. The horizontal red line indicates significance level at a false discovery rate ≤ 0.05.
